# Causal inference for heritable phenotypic risk factors using heterogeneous genetic instruments

**DOI:** 10.1101/2020.05.06.077982

**Authors:** Jingshu Wang, Qingyuan Zhao, Jack Bowden, Gibran Hemani, George Davey Smith, Dylan S. Small, Nancy R. Zhang

## Abstract

Over a decade of genome-wide association studies (GWAS) have led to the finding of extreme polygenicity of complex traits. The phenomenon that “all genes affect every complex trait” complicates Mendelian Randomization (MR) studies, where natural genetic variations are used as instruments to infer the causal effect of heritable risk factors. We reexamine the assumptions of existing MR methods and show how they need to be clarified to allow for pervasive horizontal pleiotropy and heterogeneous effect sizes. We propose a comprehensive framework GRAPPLE to analyze the causal effect of target risk factors with heterogeneous genetic instruments and identify possible pleiotropic patterns from data. By using GWAS summary statistics, GRAPPLE can efficiently use both strong and weak genetic instruments, detect the existence of multiple pleiotropic pathways, determine the causal direction and perform multivariable MR to adjust for confounding risk factors. With GRAPPLE, we analyze the effect of blood lipids, body mass index, and systolic blood pressure on 25 disease outcomes, gaining new information on their causal relationships and the potential pleiotropic pathways.

## 1 Introduction

Understanding the pathogenic mechanism of common diseases is a fundamental goal in clinical research. As randomized controlled experiments are not always feasible, researchers are looking towards Mendelian Randomization (MR) as an alternative method for probing the causal mechanisms of common diseases [1]. MR uses inherited genetic variations as instrumental variables (IV) to interrogate the causal effect of heritable risk factor(s) on the disease of interest. The basic idea is that at these variant loci, the inherited alleles are randomly transmitted from the parents to their offsprings according to Mendel’s laws. Thus, the genotypes are independent from non-heritable confounding variables which may obfuscate causal estimation in parent-offspring studies. More generally, such independence also approximately holds for population data such as those collected in genome-wide association studies (GWAS) when individuals share the same ancestry [2]. With the accumulation of data from GWAS, there is an increasing interest in MR approaches, especially in approaches that only rely on GWAS summary statistics that are publicly available [2, 3].

How well Mendelian Randomization works depends on how well the genetic variant loci used as instruments abide by the rules of IV. These rules dictate that, if the genetic locus has an effect on the disease outcome, it should be only through pathways mediated by the risk factor(s) of interest. This rule, termed exclusion restriction, is violated when there is horizontal pleiotropy, defined as the case where the genetic variant can influence the disease through pathways other than the given risk factor(s) [4]. There has been much recent attention on this issue [5–16] in MR, yet our understanding is far from complete. Current methods rely on different assumptions on the pattern of horizontal pleiotropy, often driven by statistical convenience rather than what geneticists have learned from real data. What assumptions on pleiotropy and genetic effects would be suitable? Would it be possible to learn the degree of pleiotropy from the data? Could we perform model diagnosis utilizing only GWAS summary statistics?

The pleiotropy issue that muddles Mendelian Randomization studies is, in a large part, due to the fact that complex traits are extremely polygenic [15, 17–24]. Accumulating evidence from GWAS studies indicates that many complex diseases may have an omnigenic architecture where all genes affect every complex trait [25]. While a few genes might be “core” genes, almost all genes are involved and can exert non-zero effects on diseases and their risk factors. Thus, in an MR study, many genetic instruments, if not all, may affect the disease through their effects on other unmeasured risk factors. In other words, in an MR analysis, not only would we expect horizontal pleiotropy to be a pervasive issue across all genetic variants, any disease or complex risk factor would also be associated with a large number of SNPs across the whole genome. Many existing MR methods rely on the assumption that pleiotropic effects sparsely involve only a few SNPs, which directly counters these recent insights. Methods that don’t assume sparsity often require that the pleiotropic effects cancel each other across SNPs, named as the instrument strength independent of direct effect (InSIDE) assumption [6], which can be rather optimistic. Recently, a few new methods relaxed the InSIDE assumption to consider “directional pleiotropy” through one pleiotropic pathway [11, 12, 15, 26]. However, when there exists pleiotropic pathways, there would often be an issue in identifying the true causal effect of the risk factor, and most methods are restrictive to allow for only one pleiotropy pathway. Armed with these assumptions, most existing methods also utilize only the few SNPs that have the strongest association with the risk factor as instruments, ignoring the SNPs that are weakly associated. In this work, we will show that weakly associated SNPs are also informative, and that a model combining weak and strong SNPs can increase the accuracy and stability of our estimations in some scenarios.

We propose a comprehensive statistical framework for causal effect estimation under the realistic assumption that pleiotropy may be pervasive across the genome. The framework, called GRAPPLE (Genome-wide mR Analysis under Pervasive PLEiotropy), facilitates interactive identification of multiple pleiotropic pathways and incorporates all SNPs associated with the risk factor into the analysis. GRAPPLE builds upon a previous statistical framework we developed called MR-RAPS [10] under the InSIDE assumption, but is much more comprehensive and flexible. GRAPPLE emphasizes the detection of multiple pleiotropic pathways when the InSIDE assumption is violated as well as the determination of the causal direction. GRAPPLE further addresses two common challenges in the practice: how to jointly estimate the effects with multiple risk factors to reduce directional pleiotropy, as well as how to integrate cohorts with overlapping samples. The estimation accuracy of GRAPPLE is examined through validations involving both real studies and simulations.

We apply GRAPPLE to investigate the causal effects of 5 risk factors (three plasma lipid traits, body mass index, and systolic blood pressure) on 25 common diseases. Although there have been several causal effect screening studies [9,15,26] for these risk factors and diseases, the multi-modality analysis enabled by GRAPPLE brings forth new insights on the pleiotropic landscape of these diseases and, thus, an improved understanding of the causal risk factors. Specifically, we will reexamine the role of lipid traits on coronary artery disease and type-II diabetes, where the results from MR studies have been under heated debate [2, 27, 28].

## 2 Model Overview

### 2.1 From the causal model to GWAS summary statistics

Our framework starts with a set of structural equations that jointly specify the generative model on the disease *Y* that relies on *K* observed risk factors ***X*** = (*X*_1_ …, *X_K_*) of interest, and all genetic variants ***Z*** = (*Z*_1_, *Z*_2_,…) (Fig 1a).

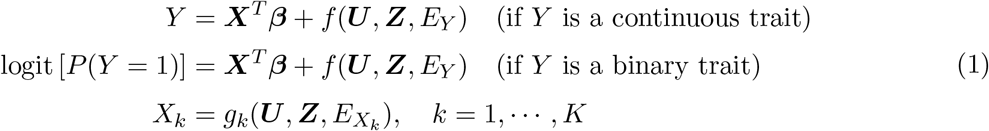

where ***U*** represents unknown non-heritable confounding factors and *E_X_k__* and *E_Y_* are random noise acting on *X_k_* and *Y* respectively. The parameter of interest, ***β*** quantifies the causal effect of the vector of risk factors ***X*** on *Y*. Mendel’s laws of inheritance suggest that the genotypes ***Z*** are randomized during conception and are generally independent of the environmental factors (***U***, *E_Y_, E_X_k__*). The function *f*(***U, Z***, *E_Y_*) represents the causal effect of unmeasured risk factors on *Y*, which can be heritable (contributed by ***Z***) or non-heritable (contributed by ***U***). The non-parametric functions *f* (·) and *g_k_*(·) allow interactions among SNPs in ***Z*** and variables (***U***, *E_Y_*, *E_X_k__*) in their causal effects on ***X*** and *Y*. Under this model, there is horizontal pleiotropy for a SNP *j* if *Z_j_* has nonzero association with *f*(***U***, ***Z***, *E_Y_*). This is the case, for example, when *Z_j_* acts on *Y* through a pathway affecting unmeasured risk factors, or when *Z_j_* is in linkage disequilibrium (LD) with such a locus.

**Fig 1.**
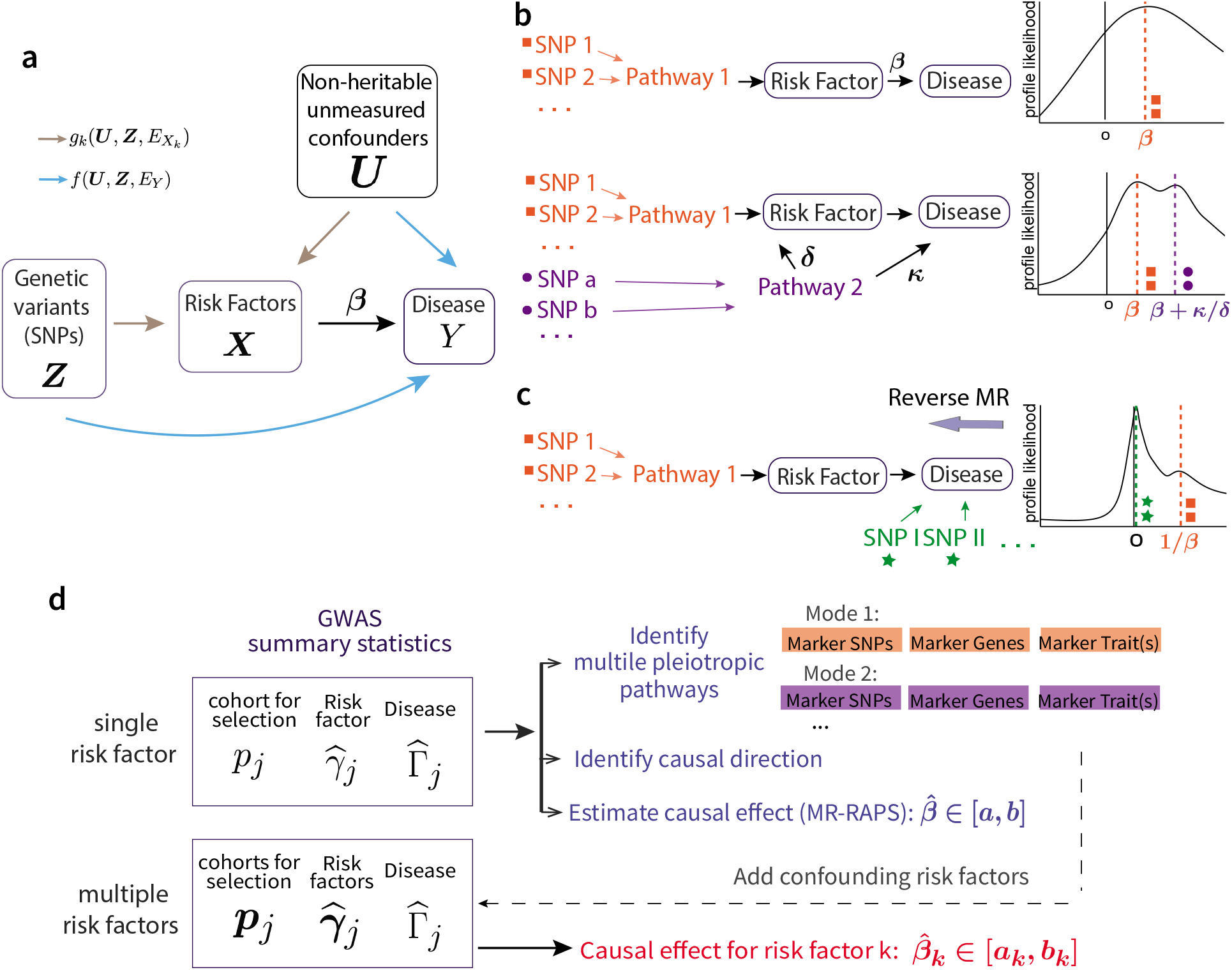
Model overview. **a**, The causal directed graph represented by structural equations (1). **b**, The existence of a pleiotropic pathway 2 (purple) can result in multiple modes of the profile likelihood. **c**, Multi-modality of the profile likelihood can reflect causal direction. **d**, The work-flow with GRAPPLE.

Now consider the case where only GWAS summary statistics, i.e. the estimated marginal associations between each SNP *j* and the risk factors/disease traits, are available and there are in total *p* SNPs selected. Let Γ_*j*_ be the true association between SNP *j* and *Y*, and ***γ***_*j*_ be the vector of true marginal associations between SNP *j* and ***X***. Later, we will denote their estimated values from GWAS summary statistics as 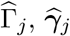. Then, as shown in Materials and Methods, the model (1) results in the linear relationship

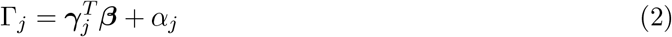

where for binary *Y*, the parameter ***β*** in (2) is a conservatively biased version of ***β*** in (1). This relationship holds even when the functions *f*(·) and *g_k_*(·) in (1) are not linear. Here, *α_j_* is the marginal association between *Z_j_* and *f*(***U***, ***Z***, *E_Y_*), representing the unknown horizontal pleiotropy of SNP *j*.

One can immediately see that identifying ***β*** is impossible without further assumptions on *α_j_*. Early MR methods such as IVW [5] made the assumption that all instruments are valid satisfying *α_j_* = 0. Other methods such as Weighted Median [7] or MR-PRSSO [9] assume that *α_j_* is sparsely nonzero. However, the no or sparse pleiotropy assumption follows from statistical convenience rather than biological insights. As discussed in Introduction, horizontal pleiotropy is pervasive for most complex traits. One assumption that allows pervasive pleiotropy is to assume the InSIDE assumption [6] where 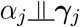, or alternatively, the random effect model [10, 16] where 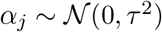 for most genetic instruments. Unfortunately, the InSIDE assumption requires all unmeasured heritable risk factors of the disease to be genetically uncorrelated with the target risk factor(s) ***X***, which is often not true.

Some more recent MR methods such as LCV [26], MRMix [12], Contamination mixture [13] and CAUSE [15] have noticed this limitation of the InSIDE assumption and allow a subset of the genetic instruments to be associated with a common hidden pleiotropic pathway. For instance, using the above notation, both CAUSE and MRMix assumed that when for the SNPs that violate the inSIDE assumption, their pleiotropic effects satisfy 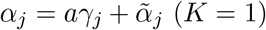 (*K* = 1) where *aγ_j_* represents the directional pleiotropic effects due to a confounding pathway and 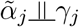. This is a more realistic assumption than InSIDE, though there would then be an issue in distinguishing the true causal effect *β* from the pleiotropic direction *β* + *a*. Allowing for only one pleiotropic pathway also makes the model restrictive for real datasets.

### 2.2 Identify multiple pleiotropic pathways and the direction of causality

The key idea underlying GRAPPLE is that multiple pleiotropic pathways can be detected by using the shape of the profile likelihood function under no pleiotropy assumption. This allows us to probe the underlying causal mechanism, without explicit assumptions of the pleiotropic patterns (Fig 1b). When *K* = 1, the GWAS summary statistics reduce to the scalar 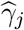 and 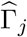,, with their standard errors *σ*_1*j*_ and *σ*_2*j*_. From the central limit theorem, the joint distribution of 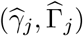 approximately follows a multivariate normal distribution

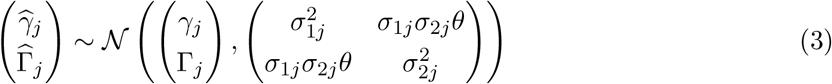

where *θ* is a shared sample correlation that can be estimated as 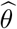 (see Materials and Methods).

When there is no horizontal pleiotropy in the *p* selected independent genetic instruments (*α_j_* = 0 for *j* = 1,2, …, *p*), the robust profile likelihood [10] is given by,

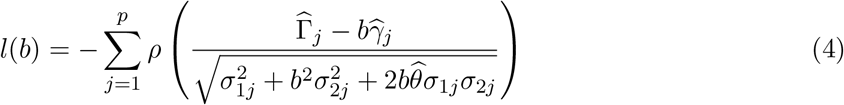

where *ρ*(·) is the Tukey’s Biweight loss, or any other robust los functions. As described with more details in Materials and Methods, the profile likelihood is obtained by profiling out nuisance parameters *γ*_1_, …, *γ_p_* in the full likelihood from (3), which is further robustified by replacing the *L*_2_ loss with Tukey’s Biweight loss to increase the sensitivity of mode detection. Under no pleiotropy or InSIDE assumption, this function *l*(*b*) should have only one mode near the true causal effect *b* = *β*.

Now consider the case where a second genetic pathway (Pathway 2) also contributes substantially to the disease, and some instrument loci are also associated with Pathway 2 (Fig 1b). In this scenario, SNPs that are associated with *X* only through Pathway 2 can contribute to a second mode in the profile likelihood at location *β* + *κ*/*δ*, where *κ* and *δ* quantifies the causal effect of Pathway 2 on *Y* and its marginal association with *X*, respectively (Materials and Methods). Similarly, multiple pleiotropic pathways generally result in multiple modes of *l*(*b*). Thus, we can use multiple modes in a plot of *l*(*b*) to diagnose the presence of horizontal pleiotropic effects that are grouped by different pleiotropic pathways.

The existence of pleiotropic pathways complicates MR and makes the causal effects of the risk factors potentially unidentifiable. Specifically, when Pathway 2 exists, the GWAS summary statistics alone can not provide information to distinguish *β* from *β* + *κ*/*δ*. Instead of making further untestable assumptions such as one pathway “dominates” the other, when multiple modes are detected, we suggest that whenever multiple modes are detected, the investigator should try to find biomarkers for each mode and collect more GWAS data to adjust for confounding risk factors. Specifically, GRAPPLE facilitates this by identifying marker SNPs of each mode, as well as the mapped genes and GWAS traits of each marker SNP (see Materials and Methods). This allows researchers to use their expert knowledge to infer possible confounding risk factors that contribute to each mode. With GWAS summary statistics of these confounding traits, GRAPPLE can perform a multivariable MR analysis assuming InSIDE assumption on the remaining horizontal pleiotropic effects (Materials and Methods).

The detection of multiple modes can be also used to determine the causal direction (Fig 1c). If the wrong causal direction is specified in (1) and *Y* is a cause of *X*, the genetic variants associated with *X* can be classified in two groups: those associated with *X* through *Y*, and those associated with *X* through another pathway unrelated to *Y*. In the former case, *γ_j_* = *β*Γ_*j*_ where *β* is the causal effect of *Y* on *X*, and these SNPs should contribute to a mode around 1/*β*. In the latter case, a SNP *j* satisfies *γ_j_* ≠ 0 but Γ_*j*_ = 0, and would contribute to a mode of *l*(*b*) at 0. Thus there will be two modes in the robust profile likelihood with one mode being around 0. This idea can be viewed as an extension to the bidirectional MR [29, 30]. Bidirectional MR is based on the assumptions that when *X* is a cause of *Y*, most of the genetic instruments for *Y* should be unassociated with *X*, because they affect *Y* through a different pathway, thus the reserve MR would indicate a zero effect on *Y* on *X*. GRAPPLE makes this inference more robust by making use of the fundamentally different shape of the robust profile likelihood plots in different directions. In the correct causal direction, the plot should only show one mode around the true causal effect *β*. In the incorrect reversed direction, the plot has two modes, one around 0 and one around 1/*β*.

### 2.3 Weak genetic instruments: A curse or a blessing?

Besides the assumption of no-horizontal-pleiotropy, for a SNP to be a valid genetic instrument, it needs to have a non-zero association with the risk factor of interest. In most MR pipelines, SNPs are selected as instruments only when their p-values are below 10^−8^, which is required to guarantee a low family-wise error rate (FWER) for GWAS data. Using such a stringent threshold also avoids weak instrument bias [31], where measurement errors in 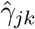 are too large to lead to bias in 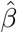. However, such a stringent selection threshold may result in very few, or even no instruments being selected with under-powered GWAS, and may still not be adequate to avoid weak instrument bias. Further, when our goal is to jointly model the effects of multiple risk factors (the setting where ***X*** as a vector), it is unrealistic to assume that all selected SNPs have strong effects on every risk factor. In addition, the high polygenecity of complex traits indicates that the weak instruments far outnumbers strong instruments, and collectively, they may substantially improve the estimation accuracy.

In GRAPPLE, we use a flexible p-value threshold, which can be either as stringent as 10^−8^ or as mild as 10^−2^, for instrument selection. Based on the profile likelihood framework of MR-RAPS [32], GRAPPLE can provide valid inference for 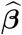 that avoids weak instrument bias for multiple risk factors even when the p-value threshold is as large as 10^−2^. This flexible p-value threshold is beneficial for several reasons. First, including moderate and weak instruments may increase power, especially for under-powered GWAS. Second, for MR with multiple risk factors where it is inevitable to include SNPs that have weak associations with some of the risk factors, we can obtain much more accurate causal effect estimations than methods that can only deal with jointly strong SNPs. More importantly, we can investigate the stability of the estimates across a series of p-value thresholds and get a more complete picture of the underlying horizontal pleiotropy. In practice, we suggest researchers to vary the selection p-value thresholds from a stringent one (say 10^−8^) to a mild one (say 10^−2^), both in the detection of multiple modes and in estimating causal effects.

### 2.4 The three-sample design to guard against instrument selection bias

The current two-sample design of MR uses one GWAS data for the risk factor and one for the disease. The selection of genetic instruments are performed with p-values reported in the GWAS data for the risk factor. However, selecting instruments from GWAS summary statistics can introduce bias, which is commonly referred to as the “winner’s curse”. Conditional on being selected, the magnitude of 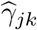 is generally larger than *γ_jk_* and introduces bias to the estimation of ***β***. When *K* = 1 where there is only one risk factor, the estimate will be biased towards 0, but there is no guarantee on the direction of the bias when *K* > 1. Among practitioners, a common belief is that the selection bias is negligible when only the strongly associated SNPs are selected as instruments.

However, this rule of thumb may not hold even when we only use that are genome-wide significant (p-value ≤ 10^−8^) (Fig S1a). Thus, we strongly advocate using a three-sample GWAS summary statistics design (Fig 1d). To avoid the selection bias, selection of genetic instruments is done on another GWAS dataset for the risk factor, whose cohort has no overlapping individuals with both the risk factor and disease cohorts. In addition, to simplify calculation and avoid bias due to different LD structure in heterogeneous populations, (see Materials and Methods), we use LD clumping [33] to select independent SNPs in GRAPPLE. The three-sample design will also avoid possible selection bias introduced during clumping.

Summarizing the above points, a complete diagram of the GRAPPLE workflow is shown in Fig 1d. A researcher may start with a single target risk factor of interest. The shape of the robust profile likelihood provides information on possible pleiotropic pathways. If only a single mode is detected, one can use GRAPPLE for the target risk factor. This is equivalent to using the original MR-RAPS. If multiple modes are detected, the researcher needs to seriously consider how to adjust for pleiotropic pathways. Researchers can use the marker SNP/gene/trait information that GRAPPLE provides to investigate each mode, decide on which confounding risk factors to adjust for, and collect extra GWAS data for them. GRAPPLE can then be used to jointly estimate the causal effects of the original and the additional risk factors.

## 3 Assessment of GRAPPLE with real studies

### 3.1 Combine weak and strong genetic instruments under no pleiotropy

We first examine whether GRAPPLE provides reliable statistical inference using instruments with different strength under an artificial setting with real GWAS summary statistics. In this setting, we make the “artificial risk factor” *X* and the “artificial disease” *Y* be the same trait from two non-overlapping cohorts, thus *γ_j_* = Γ_*j*_ while 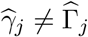 for any SNP. Though the structural equation describing the causal effect of *X* on *Y* is not well defined., the linear relationship model (2) from which we estimate *β* still holds with *β* = 1 and *α_j_* = 0. In other words, we are not estimating a meaningful “casual” effect, but are in a special case where the true *β* is known. This setup can be used to verify the validity of MR methods under no pleiotropy.

Specifically, we consider three traits: Body mass index (BMI), Type II diabetes (T2D) and height from the GIANT and DIAGRAM consortium where sex-specific GWAS data are available [34, 35]. The female cohort is used to get 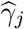 and the male cohort is used to get 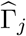. As a three-sample design, the UK Biobank data for corresponding traits are used for SNP selection. If we assume that all selected instruments have no gender-specific association with the traits, the ture *β* would equal 1. For benchmarking, we compare the performance of GRAPPLE with CAUSE [15] and other three well-adopted MR methods, inverse-variance weighted (IVW) [5], MR-Egger [6] and weighted median [7] with the same three-sample design.

We compare different p-value thresholds for instrument selection, ranging from a stringent threshold 10^−8^ to a mild threshold 10^−2^ (Fig 2a). GRAPPLE provides roughly unbiased estimates of *β* no matter which threshold is used, showing that it does not suffer from the weak instrument bias. Surprisingly, the other MR methods are biased even with a stringent p-value threshold.

**Fig 2.**
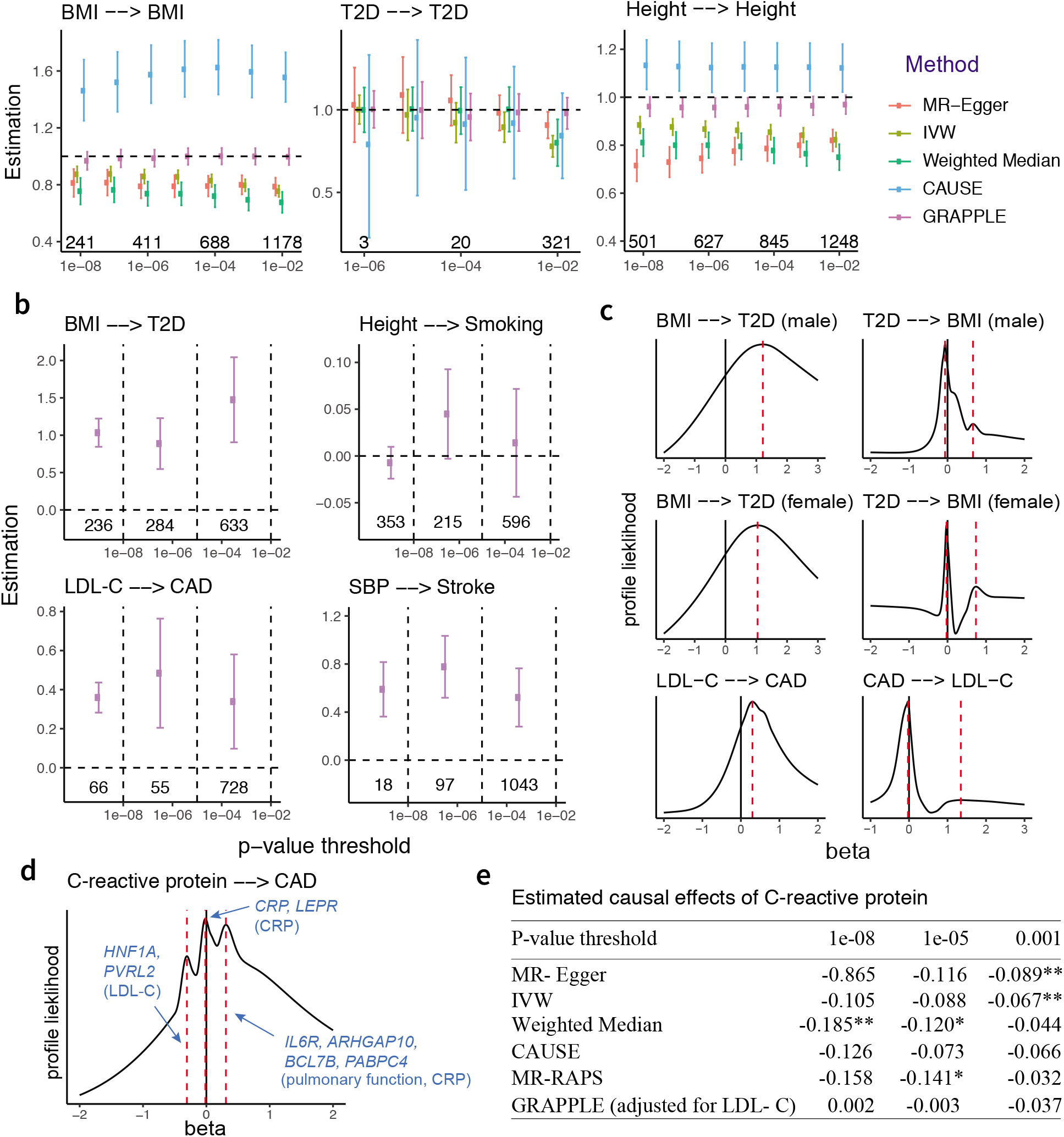
Performance evaluation. **a**, Estimation of *β* across selection p-value thresholds under no pleiotropy. Error bars show 95% Confidence intervals and the numbers are the number of independent SNPs obtained at each threshold. **b**, Estimation of *β* across three non-overlapping categories of SNPs: “strong”, “moderate” and “weak”. The numbers are the number of SNPs in each category. **c**, Identifying causal directions by multi-modality with MR reversely performed. The selection p-value threshold is kept at 10^−4^. **d**, three modes detected in the profile likelihood with selection p-value threshold 10^−4^ for CRP on CAD. Marker genes and GWAS traits (in parenthesis) are shown for each mode. **e**, estimation of the CRP effect *β* at different p-value selection threshold with each method. The numbers are the estimated 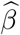, with * indicating p-value below 0.05 and ** indicating p-value below 0.01.

Notice that for T2D, the confidence intervals of GRAPPLE do get narrower with increasing p-value thresholds (Fig 2a), showing the potential power gain of including weak instruments in less powerful GWAS studies. In addition, we simulate synthetic GWAS summary statistics of the risk factor and disease (see Supplementary Note 2 for details) and confirm that the estimated *β* indeed gets more accurate with the inclusion of weakly associated SNPs (Figure S2a). In the simulations, we also use GRAPPLE to adjust for measured confounding risk factors and compare the performance with MVMR [63], a commonly used multivariable MR method. As discussed earlier, in multivariable MR, the inclusion of SNPs that are weakly associated with at least one risk factors is inevitable. As GRAPPLE do not suffer from weak instrument bias, we see that it provides accurate estimates of the causal effects as well as reliable confidence intervals with both stringent and mild p-value thresholds (Fig S3-S5).

Finally, we demonstrate that to avoid bias, the three sample design is necessary no matter which MR method is used. As shown in Fig S1a, the two-sample design where we use the same cohort of the risk factor for selection can result in biased casual effects estimation, and the bias occurs with most MR methods even when we only select the strongly associated SNPs.

### 3.2 Weak SNPs provide reliable causal estimates under pleiotropy

Next, we examine whether or not the weak instruments are more vulnerable to pleiotropy, which can be a concern for including the weak SNPs. We compare four risk factor and disease pairs that cover eight different complex traits, including the effect of BMI on T2D, low-density cholesterol concentrations (LDL-C) on coronary artery disease (CAD), height on smoking, and systolic blood pressure (SBP) on stroke (Fig 2b).

We test whether independent sets of strongly and weakly associated SNPs can provide consistent estimates of the causal effects of the risk factors. SNPs passing the p-value threshold 10^−2^ in the cohort for selection are divided into three non-overlapping groups after LD clumping: “strong” (*p_j_* ≤ 10^−8^), “moderate” (10^−8^ < *p_j_* ≤ 10^−5^), and “weak” (10^−5^ < *p_j_* ≤ 10^−2^). The SNPs across groups are used separately to obtain group-specific estimates of the causal effect *β*. We observe that for all the four pairs, the estimates 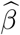 are stable across groups (Fig 2b). Though the “weaker” SNPs provide estimates with more uncertainty due to limited power, the estimates are consistent with those from the “strong” group. Other MR methods also show some level of consistency in estimating *β* across different sets of instruments, but perform worse due to weak instrument bias (Fig S1b). To conclude, in the analysis of these four pairs of traits, we do not see any evidence that weakly associated SNPs provide more biased estimates than strong instruments due to horizontal pleiotropy. In contrast, as the strong SNPs, they may also provide useful information to infer the causal effects of the risk factors.

### 3.3 Identify direction of causality for known causal relationships

We also examine the performance of GRAPPLE in identifying the causal direction with the shape of the profile likelihood. For the causal direction, we focus on the two pairs of traits with known causal relationship: BMI on T2D, and LDL-C on CAD. We switch the roles of the risk factor and disease to see if the correct direction can be revealed. Specifically, we treat T2D and CAD as the “risk factor”, and BMI and LDL-C as the corresponding “disease” (Fig 2c). For T2D, the cohort for the other gender is used for SNP selection and for CAD, the risk factor cohort used is from [36] and the selection p-values are from [37]. As expected, we see that when the roles of the risk factor and disease are reversed, the robust profile likelihood shows a main mode at 0, and a weaker mode around 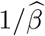.

### 3.4 Detect multiple modes to identify pleiotropic pathways

Finally, we test the ability of GRAPPLE to identify multiple pleiotropic pathways with the analysis of the C-reactive protein (CRP) effect on CAD. C-reactive protein has been found to be strongly associated with the risk of heart disease while many SNPs who are associated with the C-reactive protein also seem to have pleiotropic effect on lipid traits [38]. Previous MR analyses only included SNPs that are near the *CRP* gene to guarantee a free-of-pleiotropy analysis [39] and found that CRP has no causal effect on CAD. This is also validated by randomized experiments [40]. Now, instead of only using SNPs near the CRP gene, by using associated SNPs across the whole genome that are known to suffer from pleiotropy pathways, can GRAPPLE identify the existence of these pathways and still obtain the correct estimate of the C-reactive protein effect?

CRP GWAS data from [41] is used for selection and the data from [42] using a larger cohort is used for getting 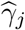. Similar to a multi-modality pattern already reported in [11], our robust profile likelihood shows a pattern of three modes, indicating the existence of at least three different pathways (Fig 2d). One mode is negative, one is positive and the third is around zero. The negative mode involves a few marker genes including *HNF1A* and *PVRL2*, with a marker trait LDL-C. The positive mode has marker traits pulmonary function and the C-reactive protein, and the few markers genes (*IL6R, ARHGAP10, BCL7B, PABPC4*) are also involved in immune response and lung cancer progression [43, 44]. The mode at 0 has marker genes *CRP* and *LEPR*, and only one marker trait, the C-reactive protein.

We compare across 3 p-value thresholds (10^−8^, 10^−5^, 10^−3^) and check how the existence of multiple pathways affects causal estimates of the effect of C-reactive protein in MR methods using SNPs across the genome. Including the C-reactive protein as the only risk factor, all bench-marking methods give a negative estimate of the CRP effect, which is possibly driven by the bias from an LDL-C induced pleiotropic pathway (Fig 2e). MR-RAPS is the estimation method used in GRAPPLE if we only use one risk factor, and the three other bench-marking methods give incorrect inference of the CRP effect with a p-value of *β* below 0.01 for at least one SNP selection threshold (notice that the weak instrument bias is towards 0 as shown in Fig 2a, thus the significance at p-value threshold 10^−3^ for MR-Egger and IVW can not be explained by weak instrument bias). In contrast, after using two risk factors: the C-reactive protein and LDL-C, where LDL-C is an identified confounding risk factor from the marker SNPs in Fig 2d, the estimates of CRP effect can keep insignificant across all p-value thresholds. In addition, the estimates 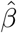 of CRP themselves are much closer to 0 compared with that without including LDL-C. This analysis illustrates how GRAPPLE can detect pleiotropic pathways, provide information to identify the confounding risk factors to adjust for, and obtain correct inference after adjusting for these risk factors.

As a complement to the analysis on CRP, we also use simulations (see Supplementary Note 2 for details) and generate synthetic disease traits to evaluate the precision and recall in the detection of multiple modes and marker SNPs when there are pleiotropic pathways. We consider scenarios with one or two pleiotropic pathways caused by hidden confounding risk factors, and vary the genetic correlations between these hidden factors and the target risk factor. A higher genetic correlation corresponds to a larger proportion of SNPs that have a directional pleiotropic effect. We observe that the detection of multiple modes is most powerful when the genetic correlation is neither too large nor too small (Fig S2b). If the genetic correlation is too high, then there are not enough SNPs to contribute to the mode of the true causal effect, while if the genetic correlation is too low there will be too few SNPs to contribute to the pleiotropic modes. Including weaker SNPs will decrease the sensitivity in mode detection but can increase the recall of true marker SNPs. In our simulations, we also observe that all univariable MR methods can perform poorly in estimating the true causal effect in the presence of pleiotropic pathways (Fig S3-S5).

## 4 A causal landscape from 5 risk factors to 25 common diseases

Finally, we apply GRAPPLE to interrogate the causal effects of 5 risk factors on 25 complex diseases. The five risk factors are three plasma lipid traits: LDL-C, high-density lipoprotein cholesterol (HDL-C), triglycerides (TG), BMI and SBP. The diseases include heart disease, Type II diabetes, kidney disease, common psychiatric disorders, inflammatory disease and cancer (Fig 3a). For each pair of the risk factor and disease, we compare across p-value thresholds from 10^−8^ to 10^−2^. As a summary of the results, Fig 3a illustrates the average number of modes detected across the p-value thresholds for SNP selection (for modes at each p-value threshold, see Figure S2). Besides the number of modes, Fig 3a also shows the p-values for each risk factor when GRAPPLE is performed with only the single risk factor (see also Fig S6, Materials and Methods). These p-values are not valid when there are pleiotropic pathways.

**Fig 3.**
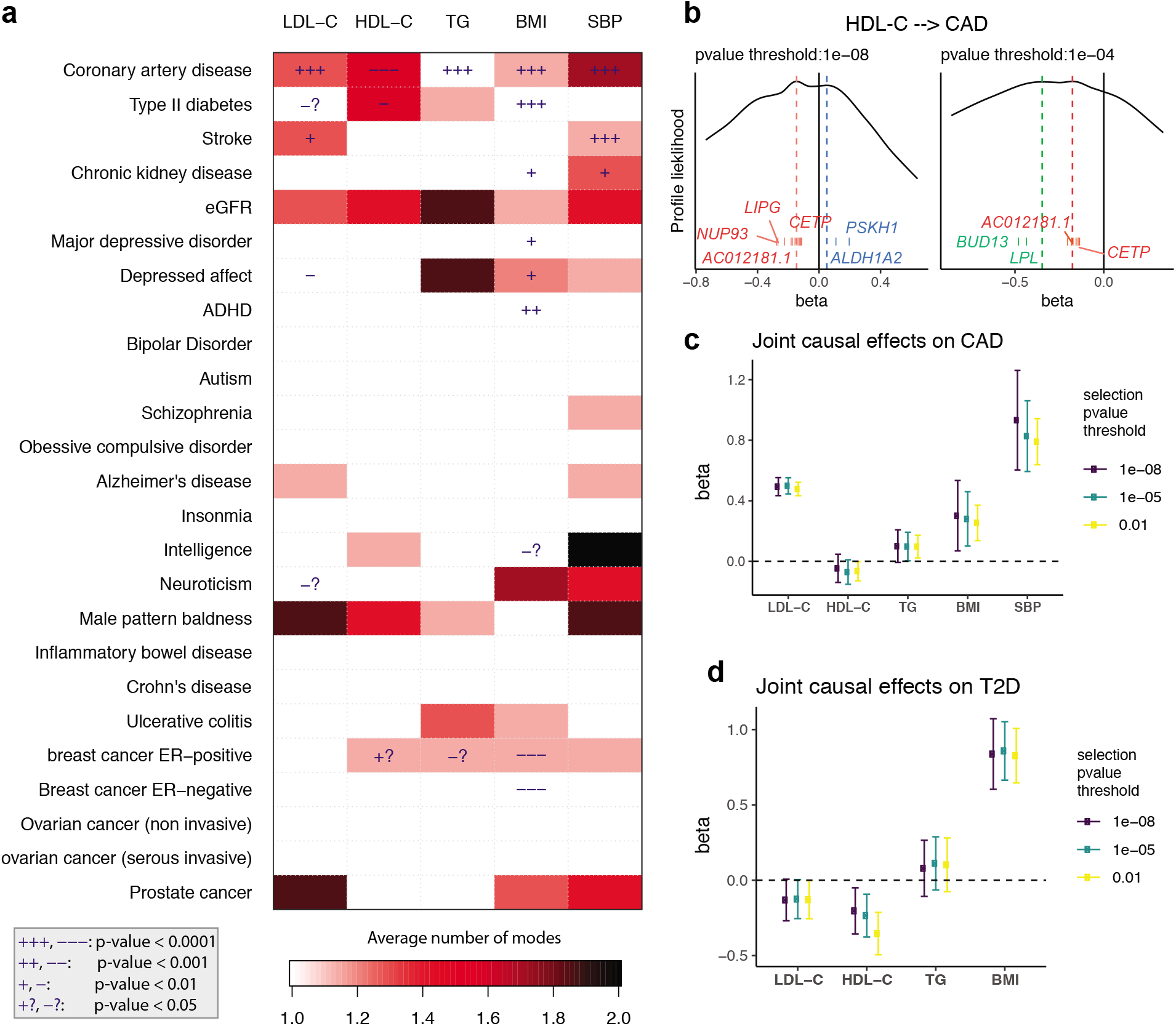
Screening with GRAPPLE. **a**, Landscape of pleiotropic pathways on 25 diseases. The colors show average number of modes across 7 different selection p-value thresholds. The “+” sign shows a positive estimated effect and “—” indicates a negative estimated effect, with the p-value for each cell a combined p-value (see Materials and Methods) of replicability across 7 thresholds using the single risk factor. These p-values are not multiple-testing adjusted across pairs. **b**, Multi-modality of the profile likelihood for effect of HDL-C on CAD at 2 different selection p-value threshold. Vertical bars are positions of marker SNPs 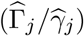, labeled by their mapped genes (only unique gene names are shown). **c**, Multivariable MR for the effect of 5 risk factors on CAD. **d**, Multivariable MR for the effect of 4 risk factors on CAD. The Error bars are 95% confidence intervals.

Fig 3a shows that multi-modality can be detected in many risk factor and disease pairs. Multimodality is most easily seen using the stringent p-value threshold 10^−8^ (Fig S6). However, we find that some modes are contributed by a single SNP thus is more likely an outlier than a pathway. For instance, the effect of stroke on LDL-C shows two modes when the p-value threshold is 10^−8^ or 10^−7^ (one mode around −2.3 and another mode near 0.08). However, the negative mode only has one marker SNP (rs3184504) which has been found strongly associated with hundreds of different traits according to GWAS Catalog [45] while the other mode has hundreds or marker genes. After removing the SNP rs3184504, the mode disappears. Such a mode also disappears when we increase the p-value threshold to include more SNPs as instruments. Thus, the average number of modes serves as a strength of evidence for the existence of multiple pleiotropic pathways. When a risk factor and disease pair show multi-modality, the p-values from GRAPPLE using the single target risk factor are no longer valid, and the researchers need further investigations of the modes.

First, consider the well-studied, often-debated relationship between CAD and the lipid traits.. All five risk factors show very significant effects, though multi-modality is detected in HDL-C and SBP. In our results for HDL-C, with different p-value thresholds, three modes in total can show up, two being negative and one positive, indicating that the pathways from HDL-C to CAD is complicated (Fig 3b). Fig 3b shows that one negative mode is contributed by SNPs near genes *LPL* and *BUD13*, which are strongly associated with triglycerides. Another positive mode is contributed by SNPs near genes *ALDH1A2* and *PSKH1*, which is related to respiratory diseases [46]. The markers of the other negative mode are mapped to genes including *LIPG* and *CETP*.

Since the effects of the lipid traits are generally complicated, we combine all 5 risk factors and run an MR jointly with GRAPPLE (Fig 3c) with different p-value thresholds. After adjusting for other risk factors, the two most prominent risk factors for the heart disease are LDL-C and SBP, while the protective effect of HDL-C stays negligible as well as the risk brought by TG. So these results show that HDL-C as a single measurement does not seem to have a protective effect on heart disease, while there are complicated multiple pathways involved. Researchers have suggested analyzing different subgroups of HDL-C as smaller particles tend to have a stronger protective effect [47].

Lipids are also involved in a number of biological functions including energy storage, signaling, and acting as structural components of cell membranes and have been reported to be associated with various diseases [48–53]. Besides CAD, another disease that most likely involves the lipid traits is the Type II diabetes (Fig 3a). T2D is associated with dyslipidemia (i.e., higher concentrations of TG and LDL-C, and lower concentrations of HDL-C), though the causal relationship is still unclear [54]. In the mean time, evidence has emerged that LDL-C reduction with statin therapy results in a modest increase in risk of T2D [48]. For the MR analyzing each risk factor alone, we see potential protective effects of LDL-C and HDL-C on T2D but also multi-modality patterns. Two modes show up in the profile likelihood from HDL-C to T2D where one negative mode has a marker gene *LPL* and a mode near 0 with marker genes *CETP* and *AC012181.1*. Thus we include all 3 lipid traits, along with BMI and run a joint model for these 4 risk factors using GRAPPLE (Fig 3d). Our result indicates a mild protective effect of HDL-C on T2D, while showing not enough evidence for the effect of either LDL-C or TG.

## 5 Discussion

We propose a comprehensive framework, GRAPPLE, that utilizes both strongly and weakly associated SNPs to understand the causal relationship between complex traits. GRAPPLE is robust to pervasive pleiotropy and can identify multiple pleiotropic pathways. The multivariable MR performed by GRAPPLE can adjust for known confounding risk factors.

GRAPPLE incorporates several improvements over existing MR methods. It gets rid of weak instrument bias by dealing with measurement errors of the SNP associations on the risk factors with profile likelihood. Our likelihood is similar to the approach in [55], while allowing pervasive pleiotropy with the InSIDE assumption. The multi-modality visualization shares similarities with [8], which estimates the causal effect by the global mode, but we provide a more comprehensive analysis to identify multiple pleiotropic pathways by the local modes. Our causality direction identification is related to bidirectional MR where they used the assumption that if we reverse the role of risk factor and disease, the estimated causal effect is likely to be 0. We use this idea in a more principled way and can avoid bias when SNPs affecting the disease through the target risk factors are also selected in the reversed MR. Finally, as the intercept term in MR-Egger is not invariant to the arbitrary assignment of effect alleles for each SNP, leading to the deficiency of the method, GRAPPLE does not include any intercept term.

GRAPPLE needs a separate GWAS cohort of the exposure for SNP selection, which is necessary for valid inference with weakly associated SNPs. Actually, as shown in Fig S1a, the three-sample design is needed for other MR methods as well to avoid selection bias. In some studies, it is hard to obtain multiple good-quality public GWAS summary statistics with non-overlapping cohorts. We call for the release of stage-specific or study-specific GWAS data summary statistics to the public in the future.

In GRAPPLE, we still require using a p-value threshold, though it can be as mild as 10^−2^, instead of requiring no p-value threshold at all. There are two main reasons for this requirement. One consideration is to increase power, as including too many SNPs with *γ_j_* = 0 or extremely small would instead increase the variance of 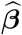 [10, 32]. Another consideration is that we would not want unmeasured risk factors that are unassociated (or very weakly associated) with target risk factors to bring in large pleiotropic effects on SNPs that mainly affect these unmeasured risk factors. The chance of including these SNPs would be much lower by requiring a mild p-value threshold.

To adjust for confounding risk factors, GRAPPLE requires that these factors are either known a priori, or can be identified from the marker SNPs / genes / traits. However, this step can be hard to execute in practice. The pleiotropic pathways may not be well tagged, and GRAPPLE may not have the power to return enough markers. As a future direction, instead of adjusting for unknown confounding risk factors, we may consider directly adjusting for confounding gene expressions that can be more easily identified.

Finally, when discussing the causal effect of a risk factor, one implicit assumption we use is consistency, assuming that there is a clear and only one version of intervention that can be done on the risk factor. However, interventions on risk factors such as BMI are typically vague [57]. For instance, there can be multiple ways to change weight, such as taking exercise, switching to different diet or conducting a surgery. It is common sense that these different interventions would have different effects on diseases, though they may change BMI by the same amount. Similarly, the cholesterol has abundant functions in our body and involves in multiple biological processes. Intervening different biological processes to change the concentrations of lipid traits may also have different effect on diseases. With MR, the interventions are changing risk factors levels with natural mutations, which may be different from interventions with drugs that has a rapid and strong effect on the risk factors. We think that our causal inference using GRAPPLE, along with the markers we detect, would provide abundant information to deepen our understanding of the risk factors. However, one still needs to be careful when giving causal interpretations of the results. One recommendation in practice is to triangulate the results from MR with other sources of evidence [58].

## Materials and methods

### Model details

The structural equations (1) where ***X*** = (*X*_1_, *X*_2_, …, *X_K_*) and ***β*** = (*β*_1_, *β*_2_, …, *β_K_*) describe how individual level data are generated. To link it with the GWAS summary statistics data, denote

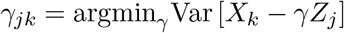

which is the true marginal association between a SNP *Z_j_* and risk factor *X_k_* and

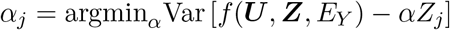

which is the marginal association between *Z_j_* and the causal effects of unmeasured risk factors on *Y*, i.e. the horizontal pleiotropic effect of *Z_j_* on *Y* given ***X***. Then we can rewrite the structural equations into the following linear models:

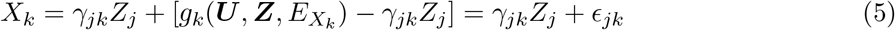

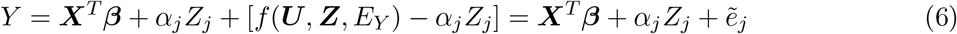

where corr(*Z_j_*, ∈_*jk*_) = 0 for any *k* and 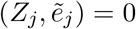 guaranteed by the definitions of *γ_jk_* and *α_j_*. By replacing ***X*** in (6) with (5), we get

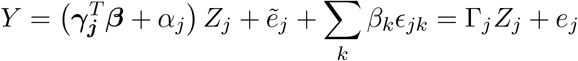

where 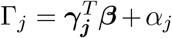 and 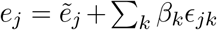 As Corr(*Z_j_*, *e_j_*) = 0, we conclude that Γ_*j*_ also satisfies that

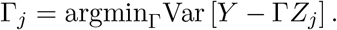

Thus, parameters Γ_*j*_ also represent true marginal associations between SNP *Z_j_* and the the disease trait. This is how we result in working with Eq (2).

When the disease is a binary trait, the structural equation of *Y* changes to

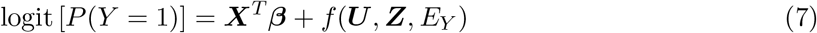

With the same argument, we have

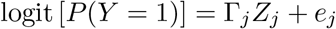

If we further assume that for each genetic instrument *j*, *Z_j_* is actually independent of *e_j_* (instead of just being uncorrelated), then the odds ratio that is estimated from the marginal logistic regression will be approximately Γ_*j*_/*c* with a constant *c* > 1 determined by the distribution of *e_j_*. In other words, for binary disease outcomes, Eq (2) is still approximately correct with the ***β*** in (2) being a conservatively biased (by a ratio of 1/*c*) version of the *β* in (7) (for a detailed calculation, see A.1 of [10]).

### GWAS summary statistics from overlapping cohorts

The GWAS estimated effect sizes (log odds ratios for binary traits) of SNP *j* are 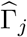 for the disease and a length *K* vector 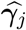 for the risk factors. As shown in [59] and derived in Supplementary Note 3.1, for any risk factor *k* we have

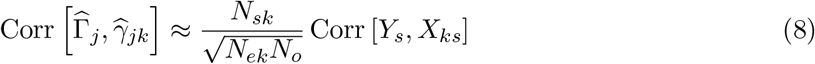

where *N_o_* and *N_ek_* are the total sample sizes for the disease and *k*th risk factor. *N_sk_* is the number of shared samples. The correlation of *X_k_* and *Y* of any shared sample is Corr [*Y_s_*, *X_ks_*]. Eq (8) shows that all the SNPs share the same correlation. As a consequence, we assume

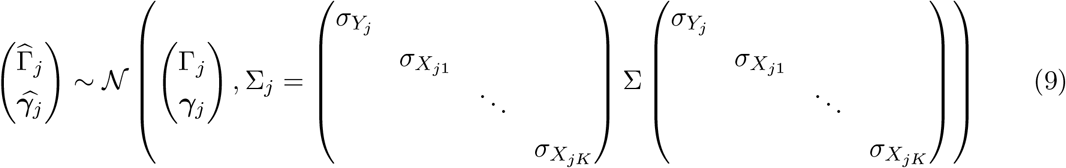

where Σ is the unknown shared correlation matrix.

### Estimate the shared correlation Σ

To estimate Σ from summary statistics, we can use Eq (8). We first need to choose SNPs where *γ_jk_* = 0 for all risk factors *k* so that we can estimate the shared correlation Corr 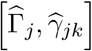 using the sample correlation of the chosen SNPs. We choose all SNPs whose selection p-values *p_jk_* ≥ 0.5 for all *k*.

For these selected SNPs, denote the Z-values of 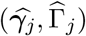 for *j* = 1, …, *T* as matrix *Z*_*T*×(*K*+1)_ where *T* is the number of selected SNPs. Then Σ is estimated as the correlation matrix of *Z*_*T*×(*K*+1)_.

### Instruments selection using LD clumping

In GRAPPLE, we need to first select a set of SNPs as genetic instruments to estimate the causal effects ***β***. Here, we only select independent SNPs to simplify the calculation. Besides the independence requirement, we only include SNPs that pass a p-value threshold to reduce the inclusion of false positives that can decrease power. To avoid selection bias, a separate cohort for each risk factor is used where the reported p-values in that cohort are used for instruments selection. Denote the selection p-value for SNP *j* and risk factor *k* as *p_jk_*, for multiple risk factors and a given selection threshold, we require the Bonferroni combined p-values *K* min(*p_jk_*) to pass the threshold. After that, we use LD clumping with PLINK [60] to select independent genetic instruments. The LD *r*^2^ threshold for PLINK is set to 0.001.

### Estimate the effects β

Here, we perform statistical analysis assuming *α_j_* ~ *N*(0, *τ*^2^) for the pleiotropic effects, while robust to outliers where the pleiotropic effects for a few instruments are large.

Under model (9), Eq (2) and given Σ, the log-likelihood with GWAS summary statistics satisfy:

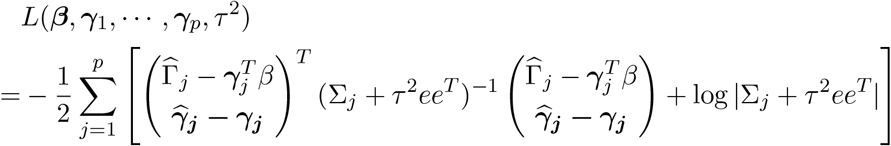

up to some additive constant. Here, *e* = (1, 0, …, 0).

Define for each SNP *j* the statistics

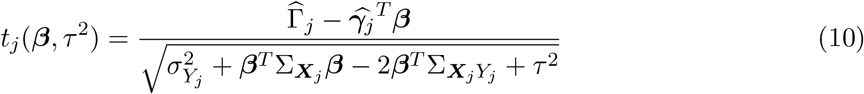

where **Σ**_*X*_*j*__ is the variance of 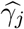 and Σ_***X***_*j*_*Y*_*j*__ is the covariance between 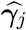 and 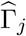 in Σ_*j*_. Then the profile log-likelihood that profile out parameters (***γ***_1_, …, ***γ***_*p*_) results in

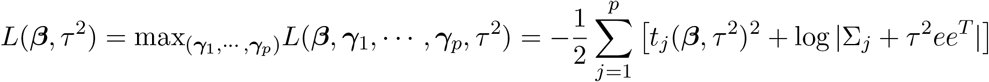

As discussed in [10], maximizing ***L***(***β***, *τ*^2^) would not give consistent estimate of *τ*^2^. Because of this and the goal of making 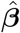 robust to outlier SNPs with large pleiotropic effects, our optimization function is the adjusted robust profile likelihood defined as

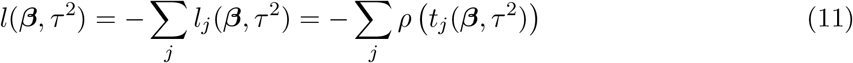

where *ρ*(·) is some robust loss function. By default, GRAPPLE uses the Tukey’s Biweight loss function:

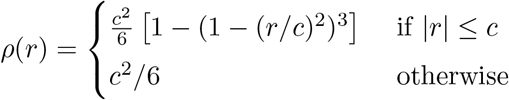

where *c* is set to its common default value 4.6851. We maximize (11) with respect to ***β*** as well as solving the following estimating equation for the heterogeneity *τ*^2^ which is

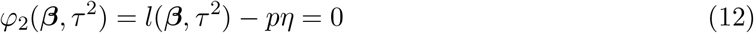

where 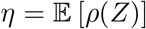 with 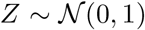. The estimating equation satisfies 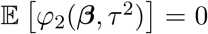 at the true values of ***β*** and *τ*^2^, thus can result in consistent estimate of *τ*^2^. For the details of estimating ***β*** and *τ*^2^ as well building confidence intervals for them, see Supplementary Note 3.2.

### Identify pleiotropic pathways via the multi-modality diagnosis

We use the mode detection of the robust profile likelihood (11) to detect multiple pleiotropic pathways. To increase sensitivity, we set *τ*^2^ = 0 and reduce the tuning parameter in the Tukey’s Biweight loss function to *c* = 3. Here we present a detailed argument on why mode detection can identify pleiotropic pathways.

If there is a confounding Genetic Pathway 2 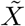, as shown in Figure 1a, that are missed, then we have the structural equation

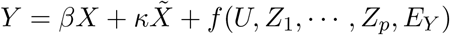

and also the linear model

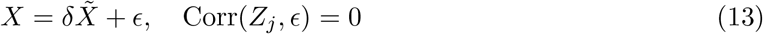

for a SNP *j* that only associate with Genetic Pathway 2 and uncorrelate with *X* conditional on 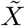. Similar to (5), we have

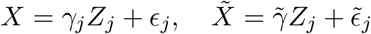

Plug in (13), we have

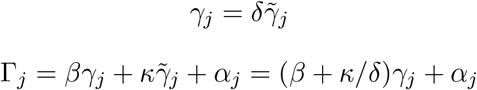

Thus, if there are enough SNPs like SNP *j*, they would contribute to another mode of (4) at *β* + *κ*/*δ*.

The same argument works for identification of the causal direction. Say there is another 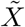 that affects *Y* but is uncorrelated with the risk factor *X* (*δ* = 0). The existence of such 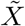 is common, unless *X* is the only heritable risk factor of *Y*. SNPs strongly associated with 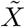 would not likely be selected when *X* is the exposure while would appear when the roles of *X* and *Y* are switched. These SNPs can be used to identify the causal direction, as as in the reverse MR, they contribute to a mode at 0, while the SNPs that affect *Y* through *X* will contribute to a mode at 1/*β*.

### Select marker SNPs and genes for each mode

GRAPPLE uses LD clumping with a stringent *r*^2^ (= 0.001) threshold to guarantee independence among the genetic instruments. However, marker SNPs are not restricted to these independent instruments in order to get more biological meaningful markers. Marker SNPs are selected from a SNP set 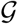 where the SNPs are selected using LD clumping with *r*^2^ threshold 0.05.

Assume that there are *M* modes detected at positions *β*_1_, *β*_2_, …, *β_M_*. Define the residual of SNP 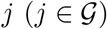 for mode *m* as

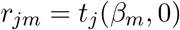

where *t_j_*(·, ·) is defined in Eq (10). SNP *j* is selected as a marker for mode *m* if |*r_jm_*’| > *t*_1_ for any *m*’ ≠ *m* and |*r_jm_*| ≤ *t*_0_. By default, *t*_1_ is set to 2 and *t*_0_ is set to 1 which gives reasonable results in practice. When the marker SNPs are selected, GRAPPLE further map the SNPs to ENCODE genes where the marker SNPs locate and and search for the traits that these SNPs are strongly associated with in GWA studies by querying HaploReg v4.1 [61] using the R package *HaploR*. The ratios 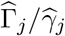 of the marker SNPs are also returned for reference (shown as the vertical bars in Fig 3b).

### Compute replicability p-values across SNP selection thresholds

Each p-value shown in Fig 3a summarizes a vector of p-values across 7 different selection p-value thresholds ranging from 10^−8^ ot 10^−2^ for each risk factor and disease pair. It reflects how consistent the significance is across SNP selection thresholds. Specifically, it is the partial conjunction p-value [62] for rejecting the null that *β* is non-zero for at most 2 of the selection thresholds. For a risk factor and disease pair *k*, let the p-values computed by using SNPs selected with the 7 thresholds *p_ks_* where s = 1, 2, …, 7. Then rank them as *p*_*k*(1)_ ≤ *p*_*k*(2)_ ≤ … ≤ *p*_*k*(7)_, the partial conjunction p-value for the pair *k* is computed as 5_*p*_*k(3)*__.

## Data and code availability

All GWAS summary statistics that are used in the analyses of the manuscript are downloaded from public resources, where most of them are downloaded from the GWAS Catalog [45], and the websites of GWAS consortium GIANT, DIAGRAM, PGC, GLGC, and UKBiobank. A complete list of the datasets used in each analysis and where they are from is provided in Supplementary Tables 1 and 2 and Supplementary Note 4. Intermediate results for screening of 5 risk factors on 25 diseases are available at https://www.dropbox.com/sh/myh8xgxne8fo17v/AABWJf781VrCGnqNFMLtnqIea?dl=0.

The R package GRAPPLE can be installed from Github at https://github.com/jingshuw/GRAPPLE.

## Supporting information

**S1 Fig. Additional evaluation results with real data. a**, Selection bias in MR methods when SNP selection and 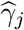 are obtained from the same GWAS dataset. True *β* ≈ 1 and error bars show 95% confidence intervals. The numbers are the number of clumped SNPs at different threshold. **b**, The estimate of *β* across three independent categories of SNPs with different association strengths for four risk factor and disease pairs using three other bench-marking MR methods. The numbers are the number of SNPs in each category, separated by the values of their selection p-values (dashed vertical lines).

**S2 Fig. Simulation results. a**, Boxplots of the estimated *β*_1_ using different MR methods over 100 repeated random experiments when there are no directional pleiotropy. We compare across three different *β*_1_ values (0.2, 0.5 and 1) with SNPs selected by three different selection thresholds: 10^−8^ for the top 106 SNPs, 10^−5^ for the top 217 SNPs and 0.01 for the top 422 SNPs. **b**, Performance of GRAPPLE in detecting multi-modality. In each setting with pleiotropic pathways, we evaluate three metrics: the detection rate of multi-modality, the precision of the identified marker genes of the pleiotropic pathways and the recall of true marker genes that are identified. Each color represent a different metric and each shape is for a different selection threshold. The title of each plot shows (*β*_1_, …, *β_K_*) in each setting where *β*_1_ is the true causal effect, and in each setting, we vary the genetic correlation between each genetic confounding risk factor and the risk factor of interest.

**S3 Fig. Comparison of different MR methods in settings with pleiotropic pathways when the selection threshold is** 10^−8^ **(top 106 SNPs). a**, Boxplots of the estimated *β*_1_ using different MR methods over 100 repeated random experiments. **b** The actual coverage of the 95% confidence intervals of *β*_1_ provided by different methods. For CAUSE, we report the coverage of the 95% credible intervals of *β*_1_. The red dotted line shows the expected 0.95 nominal level. For the three settings in the second row with *β*_1_ = 0, the CI coverage is the same as 1 — type I error.

**S4 Fig. Same as Fig. S3 with the selection threshold being** 10^−5^ **(top** 217 **SNPs).**

**S5 Fig. Same as Fig. S3 with the selection threshold being** 10^−2^ **(top** 422 **SNPs).**

**S6 Fig. Additional results on identifying pleiotropic pathways on** 25 **diseases.** Each figure is for results obtained using one of the 7 p-value thresholds. The colors show the number of detected modes. The “+” sign shows a positive estimated effect and “–” sign shows a negative estimated effect.

**S1 Table Names of the datasets used in validation analyses and screening risk factors**

**S2 Table Names of the datasets used for the** 25 **disesases in the screening application**

**Supplementary notes** Supplementary notes on the simulation details and additional mathematical details of our statistical model.

## References

1. Davey Smith G, Ebrahim S. ‘Mendelian randomization’: can genetic epidemiology contribute to understanding environmental determinants of disease? International journal of epidemiology. 2003;32(1):1–22.

2. Smith GD, Holmes MV, Davies NM, Ebrahim S. Mendel’s laws, Mendelian randomization and causal inference in observational data: substantive and nomenclatural issues. European Journal of Epidemiology. 2020; p. 1–13.

3. Davey Smith G, Hemani G. Mendelian randomization: genetic anchors for causal inference in epidemiological studies. Human molecular genetics. 2014;23(R1):R89–R98.

4. Ebrahim S, Smith GD. Mendelian randomization: can genetic epidemiology help redress the failures of observational epidemiology? Human genetics. 2008;123(1):15–33.

5. Burgess S, Butterworth A, Thompson SG. Mendelian randomization analysis with multiple genetic variants using summarized data. Genetic epidemiology. 2013;37(7):658–665.

6. Bowden J, Davey Smith G, Burgess S. Mendelian randomization with invalid instruments: effect estimation and bias detection through Egger regression. International journal of epidemiology. 2015;44(2):512–525.

7. Bowden J, Davey Smith G, Haycock PC, Burgess S. Consistent estimation in Mendelian randomization with some invalid instruments using a weighted median estimator. Genetic epidemiology. 2016;40(4):304–314.

8. Hartwig FP, Davey Smith G, Bowden J. Robust inference in summary data Mendelian randomization via the zero modal pleiotropy assumption. International journal of epidemiology. 2017;46(6):1985–1998.

9. Verbanck M, Chen Cy, Neale B, Do R. Detection of widespread horizontal pleiotropy in causal relationships inferred from Mendelian randomization between complex traits and diseases. Nature genetics. 2018;50(5):693–698.

10. Zhao Q, Wang J, Hemani G, Bowden J, Small DS, et al. Statistical inference in two-sample summary-data Mendelian randomization using robust adjusted profile score. Annals of Statistics. 2020;48(3):1742–1769.

11. Burgess S, Zuber V, Gkatzionis A, Foley CN. Modal-based estimation via heterogeneity-penalized weighting: model averaging for consistent and efficient estimation in Mendelian randomization when a plurality of candidate instruments are valid, International journal of epidemiology. 2018;47(4):1242–1254.

12. Qi G, Chatterjee N. Mendelian randomization analysis using mixture models for robust and efficient estimation of causal effects. Nature Communications. 2019;10(1):1–10.

13. Burgess S, Foley CN, Allara E, Staley JR, Howson JM. A robust and efficient method for Mendelian randomization with hundreds of genetic variants. Nature Communications. 2020;11(1):1–11.

14. Berzuini C, Guo H, Burgess S, Bernardinelli L. A Bayesian approach to Mendelian randomization with multiple pleiotropic variants. Biostatistics. 2020;21(1):86–101.

15. Morrison J, Knoblauch N, Marcus JH, Stephens M, He X. Mendelian randomization accounting for correlated and uncorrelated pleiotropic effects using genome-wide summary statistics. Nature Genetics. 2020; p. 1–7.

16. Sanderson E, Spiller W, Bowden J. Testing and Correcting for Weak and Pleiotropic Instruments in Two-Sample Multivariable Mendelian Randomisation. bioRxiv. 2020;.

17. Consortium IS. Common polygenic variation contributes to risk of schizophrenia that overlaps with bipolar disorder. Nature. 2009;460(7256):748.

18. Yang J, Benyamin B, McEvoy BP, Gordon S, Henders AK, Nyholt DR, et al. Common SNPs explain a large proportion of the heritability for human height. Nature Genetics. 2010;42(7):565–569. doi:10.1038/ng.608.

19. Bulik-Sullivan BK, Loh PR, Finucane HK, Ripke S, Yang J, Patterson N, et al. LD Score regression distinguishes confounding from polygenicity in genome-wide association studies. Nature genetics. 2015;47(3):291.

20. Loh PR, Bhatia G, Gusev A, Finucane HK, Bulik-Sullivan BK, Pollack SJ, et al. Contrasting genetic architectures of schizophrenia and other complex diseases using fast variance-components analysis. Nature Genetics. 2015;47(12):1385–1392. doi:10.1038/ng.3431.

21. Shi H, Kichaev G, Pasaniuc B. Contrasting the genetic architecture of 30 complex traits from summary association data. The American Journal of Human Genetics. 2016;99(1):139–153.

22. Timpson NJ, Greenwood CM, Soranzo N, Lawson DJ, Richards JB. Genetic architecture: the shape of the genetic contribution to human traits and disease. Nature Reviews Genetics. 2018;19(2):110.

23. O’Connor LJ, Schoech AP, Hormozdiari F, Gazal S, Patterson N, Price AL. Extreme polygenicity of complex traits is explained by negative selection. The American Journal of Human Genetics. 2019;105(3):456–476.

24. Wray NR, Wijmenga C, Sullivan PF, Yang J, Visscher PM. Common disease is more complex than implied by the core gene omnigenic model. Cell. 2018;173(7):1573–1580.

25. Boyle EA, Li YI, Pritchard JK. An expanded view of complex traits: from polygenic to omnigenic. Cell. 2017;169(7):1177–1186.

26. O’Connor LJ, Price AL. Distinguishing genetic correlation from causation across 52 diseases and complex traits. Nature genetics. 2018;50(12):1728–1734.

27. Krauss RM. Lipids and lipoproteins in patients with type 2 diabetes. Diabetes care. 2004;27(6):1496–1504.

28. Lotta LA, Sharp SJ, Burgess S, Perry JR, Stewart ID, Willems SM, et al. Association between low-density lipoprotein cholesterol–lowering genetic variants and risk of type 2 diabetes: a meta-analysis. Jama. 2016;316(13):1383–1391.

29. Timpson NJ, Nordestgaard BG, Harbord RM, Zacho J, Frayling TM, Tybjærg-Hansen A, et al. C-reactive protein levels and body mass index: elucidating direction of causation through reciprocal Mendelian randomization. International journal of obesity. 2011;35(2):300–308.

30. Hemani G, Tilling K, Smith GD. Orienting the causal relationship between imprecisely measured traits using GWAS summary data. PLoS genetics. 2017;13(11):e1007081.

31. Burgess S, Thompson SG, Collaboration CCG. Avoiding bias from weak instruments in Mendelian randomization studies. International journal of epidemiology. 2011;40(3):755–764.

32. Zhao Q, Chen Y, Wang J, Small DS. Powerful three-sample genome-wide design and robust statistical inference in summary-data Mendelian randomization. International Journal of Epidemiology. 2019;48(5):1478–1492. doi:10.1093/ije/dyz142.

33. Purcell S, Neale B, Todd-Brown K, Thomas L, Ferreira MA, Bender D, et al. PLINK: a tool set for whole-genome association and population-based linkage analyses. The American journal of human genetics. 2007;81(3):559–575.

34. Justice AE, Winkler TW, Feitosa MF, Graff M, Fisher VA, Young K, et al. Genome-wide meta-analysis of 241,258 adults accounting for smoking behaviour identifies novel loci for obesity traits. Nature communications. 2017;8:14977.

35. Morris AP, Voight BF, Teslovich TM, Ferreira T, Segre AV, Steinthorsdottir V, et al. Large-scale association analysis provides insights into the genetic architecture and pathophysiology of type 2 diabetes. Nature genetics. 2012;44(9):981.

36. Coronary Artery Disease (C4D) Genetics Consortium, et al. A genome-wide association study in Europeans and South Asians identifies five new loci for coronary artery disease. Nature genetics. 2011;43(4):339.

37. Schunkert H, König IR, Kathiresan S, Reilly MP, Assimes TL, Holm H, et al. Large-scale association analysis identifies 13 new susceptibility loci for coronary artery disease. Nature genetics. 2011;43(4):333–338.

38. Elliott P, Chambers JC, Zhang W, Clarke R, Hopewell JC, Peden JF, et al. Genetic loci associated with C-reactive protein levels and risk of coronary heart disease. Jama. 2009;302(1):37–48.

39. C Reactive Protein Coronary Heart Disease Genetics Collaboration, et al. Association between C reactive protein and coronary heart disease: Mendelian randomization analysis based on individual participant data. Bmj. 2011;342:d548.

40. Holmes MV, Ala-Korpela M, Smith GD. Mendelian randomization in cardiometabolic disease: challenges in evaluating causality. Nature Reviews Cardiology. 2017;14(10):577.

41. Prins BP, Kuchenbaecker KB, Bao Y, Smart M, Zabaneh D, Fatemifar G, et al. Genome-wide analysis of health-related biomarkers in the UK Household Longitudinal Study reveals novel associations. Scientific reports. 2017;7(1):1–9.

42. Dehghan A, Dupuis J, Barbalic M, Bis JC, Eiriksdottir G, Lu C, et al. Meta-Analysis of Genome-Wide Association Studies in¿ 80 000 Subjects Identifies Multiple Loci for C-Reactive Protein LevelsClinical Perspective. Circulation. 2011;123(7):731–738.

43. Spencer S, Köstel Bal S, Egner W, Lango Allen H, Raza SI, Ma CA, et al. Loss of the interleukin-6 receptor causes immunodeficiency, atopy, and abnormal inflammatory responses. Journal of Experimental Medicine. 2019;216(9):1986–1998.

44. Teng JP, Yang ZY, Zhu YM, Ni D, Zhu ZJ, Li XQ. The roles of ARHGAP10 in the proliferation, migration and invasion of lung cancer cells. Oncology letters. 2017;14(4):4613–4618.

45. Buniello A, MacArthur JAL, Cerezo M, Harris LW, Hayhurst J, Malangone C, et al. The NHGRI-EBI GWAS Catalog of published genome-wide association studies, targeted arrays and summary statistics 2019. Nucleic acids research. 2019;47(D1):D1005–D1012.

46. Wang J, Li F, Wei H, Lian ZX, Sun R, Tian Z. Respiratory influenza virus infection induces intestinal immune injury via microbiota-mediated Th17 cell–dependent inflammation. Journal of Experimental Medicine. 2014;211(12):2397–2410.

47. Zhao Q, Wang J, Miao Z, Zhang N, Hennessy S, Small DS, et al. The role of lipoprotein subfractions in coronary artery disease: A Mendelian randomization study. bioRxiv. 2019; p. 691089.

48. White J, Swerdlow DI, Preiss D, Fairhurst-Hunter Z, Keating BJ, Asselbergs FW, et al. Association of lipid fractions with risks for coronary artery disease and diabetes. JAMA cardiology. 2016;1(6):692–699.

49. Fonseca ACR, Resende R, Oliveira CR, Pereira CM. Cholesterol and statins in Alzheimer’s disease: current controversies. Experimental neurology. 2010;223(2):282–293.

50. Yadav RS, Tiwari NK. Lipid integration in neurodegeneration: an overview of Alzheimer’s disease. Molecular neurobiology. 2014;50(1):168–176.

51. Hibbeln JR, Salem Jr N. Dietary polyunsaturated fatty acids and depression: when cholesterol does not satisfy. The American journal of clinical nutrition. 1995;62(1):1–9.

52. Maes M, Smith R, Christophe A, Vandoolaeghe E, Gastel AV, Neels H, et al. Lower serum high-density lipoprotein cholesterol (HDL-C) in major depression and in depressed men with serious suicidal attempts: relationship with immune-inflammatory markers. Acta Psychiatrica Scandinavica. 1997;95(3):212–221.

53. Agouridis AP, Elisaf M, Milionis HJ. An overview of lipid abnormalities in patients with inflammatory bowel disease. Annals of Gastroenterology: Quarterly Publication of the Hellenic Society of Gastroenterology. 2011;24(3):181.

54. Fall T, Xie W, Poon W, Yaghootkar H, Mägi R, Knowles JW, et al. Using genetic variants to assess the relationship between circulating lipids and type 2 diabetes. Diabetes. 2015; p. db141710.

55. Burgess S, Thompson SG. Multivariable Mendelian randomization: the use of pleiotropic genetic variants to estimate causal effects. American journal of epidemiology. 2015;181(4):251–260.

56. Vimaleswaran KS, Berry DJ, Lu C, Tikkanen E, Pilz S, Hiraki LT, et al. Causal relationship between obesity and vitamin D status: bi-directional Mendelian randomization analysis of multiple cohorts. PLoS medicine. 2013;10(2):e1001383.

57. Cole SR, Frangakis CE. The consistency statement in causal inference: a definition or an assumption? Epidemiology. 2009;20(1):3–5.

58. Munafò MR, Smith GD. Robust research needs many lines of evidence; 2018.

59. Bulik-Sullivan B, Finucane HK, Anttila V, Gusev A, Day FR, Loh PR, et al. An atlas of genetic correlations across human diseases and traits. Nature genetics. 2015;47(11):1236.

60. International Schizophrenia Consortium, Purcell SM, Wray NR, Stone JL, Visscher PM, O’Donovan MC, et al. Common polygenic variation contributes to risk of schizophrenia and bipolar disorder. Nature. 2009;460(7256):748–752. doi:10.1038/nature08185.

61. Ward LD, Kellis M. HaploReg: a resource for exploring chromatin states, conservation, and regulatory motif alterations within sets of genetically linked variants. Nucleic acids research. 2012;40(D1):D930–D934.

62. Benjamini Y, Heller R. Screening for partial conjunction hypotheses. Biometrics. 2008;64(4):1215–1222.

63. Eleanor S, George DS, Frank W. and Jack B. An examination of multivariable Mendelian randomization in the single-sample and two-sample summary data settings International journal of epidemiology. 2019;48(3):713–727.

