## Supplementary tables, figures and text for "Causal inference for heritable phenotypic risk factors using heterogeneous genetic instruments"

### 1 Supplementary figures

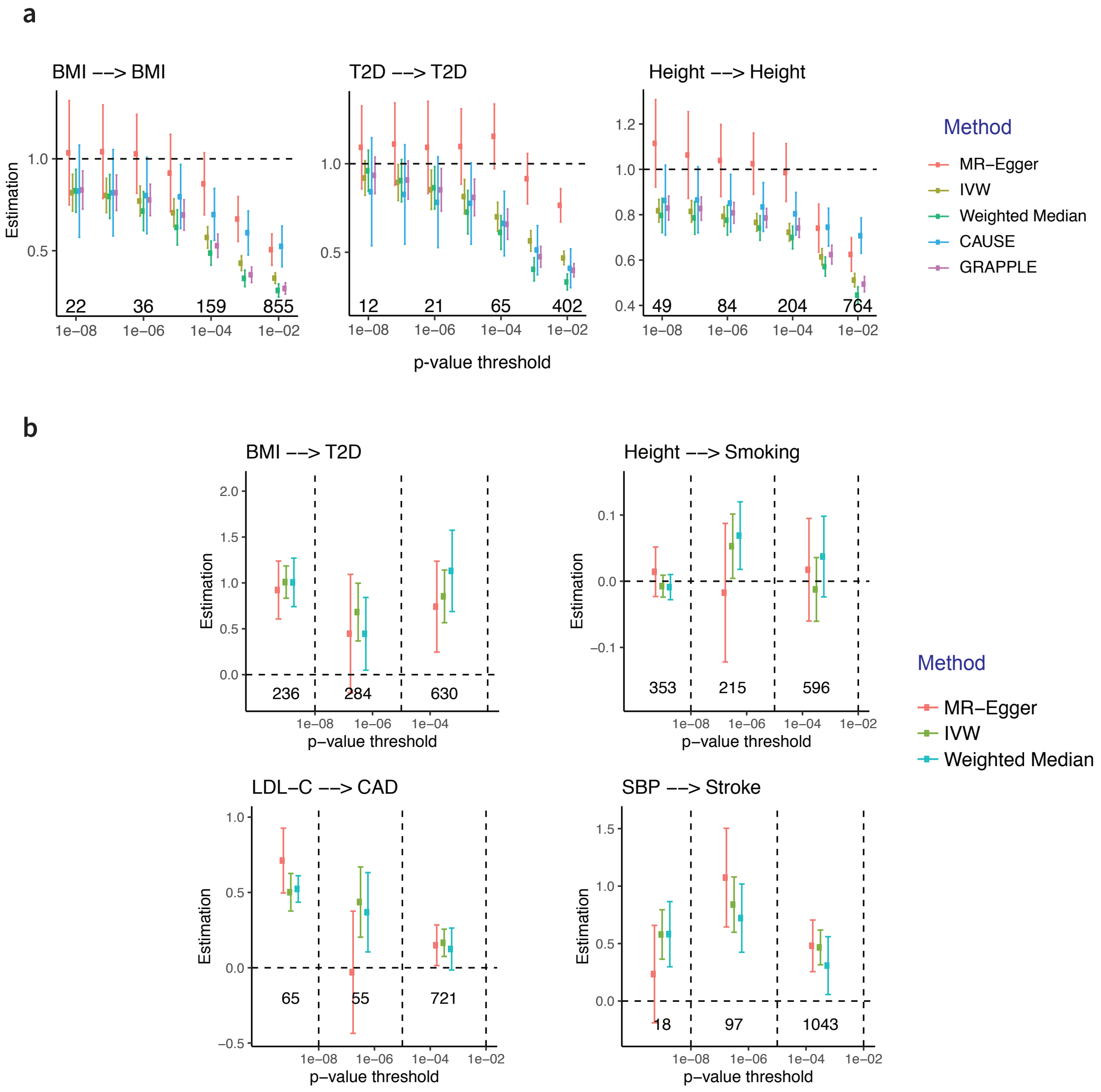


Figure S1: Additional evaluation results with real data. **a**, Selection bias in MR methods when SNP selection and $\widehat{\gamma}_j$ are obtained from the same GWAS dataset. True $\beta \approx 1$ and error bars show 95% confidence intervals. The numbers are the number of clumped SNPs at different threshold. **b**, The estimate of $\beta$ across three independent categories of SNPs with different association strengths for four risk factor and disease pairs using three other bench-marking MR methods. The numbers are the number of SNPs in each category, separated by the values of their selection p-values (dashed vertical lines).

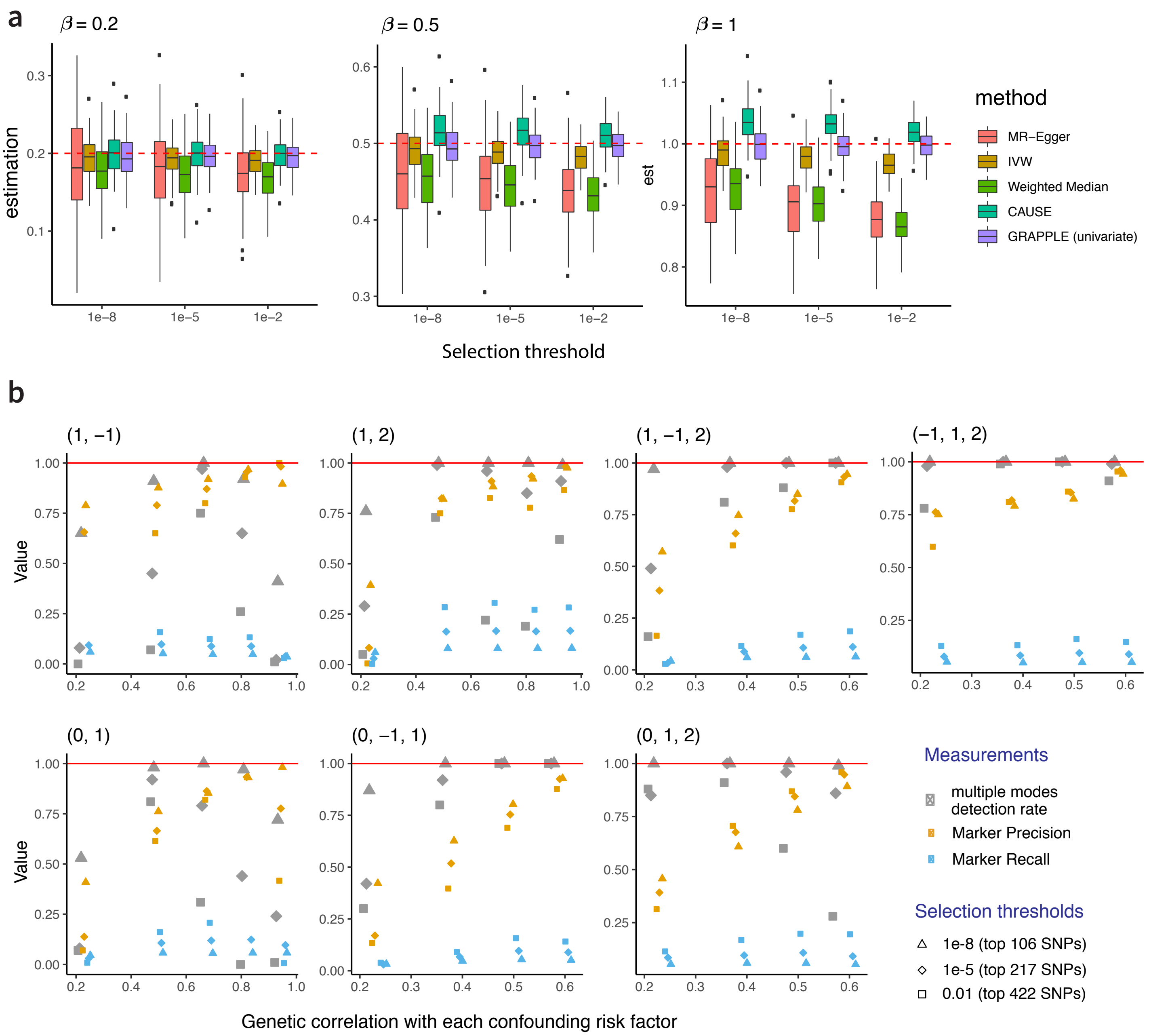


Figure S2: Simulation results. **a**, Boxplots of the estimated $\beta_1$ using different MR methods over 100 repeated random experiments when there are no directional pleiotropy. We compare across three different $\beta_1$ values (0.2, 0.5 and 1) with SNPs selected by three different selection thresholds: $10^{-8}$ for the top 106 SNPs, $10^{-5}$ for the top 217 SNPs and 0.01 for the top 422 SNPs. **b**, Performance of GRAPPLE in detecting multi-modality. In each setting with pleiotropic pathways, we evaluate three metrics: the detection rate of multi-modality, the precision of the identified marker genes of the pleiotropic pathways and the recall of true marker genes that are identified. Each color represent a different metric and each shape is for a different selection threshold. The title of each plot shows $(\beta_1, \cdots, \beta_K)$ in each setting where $\beta_1$ is the true causal effect, and in each setting, we vary the genetic correlation between each genetic confounding risk factor and the risk factor of interest.

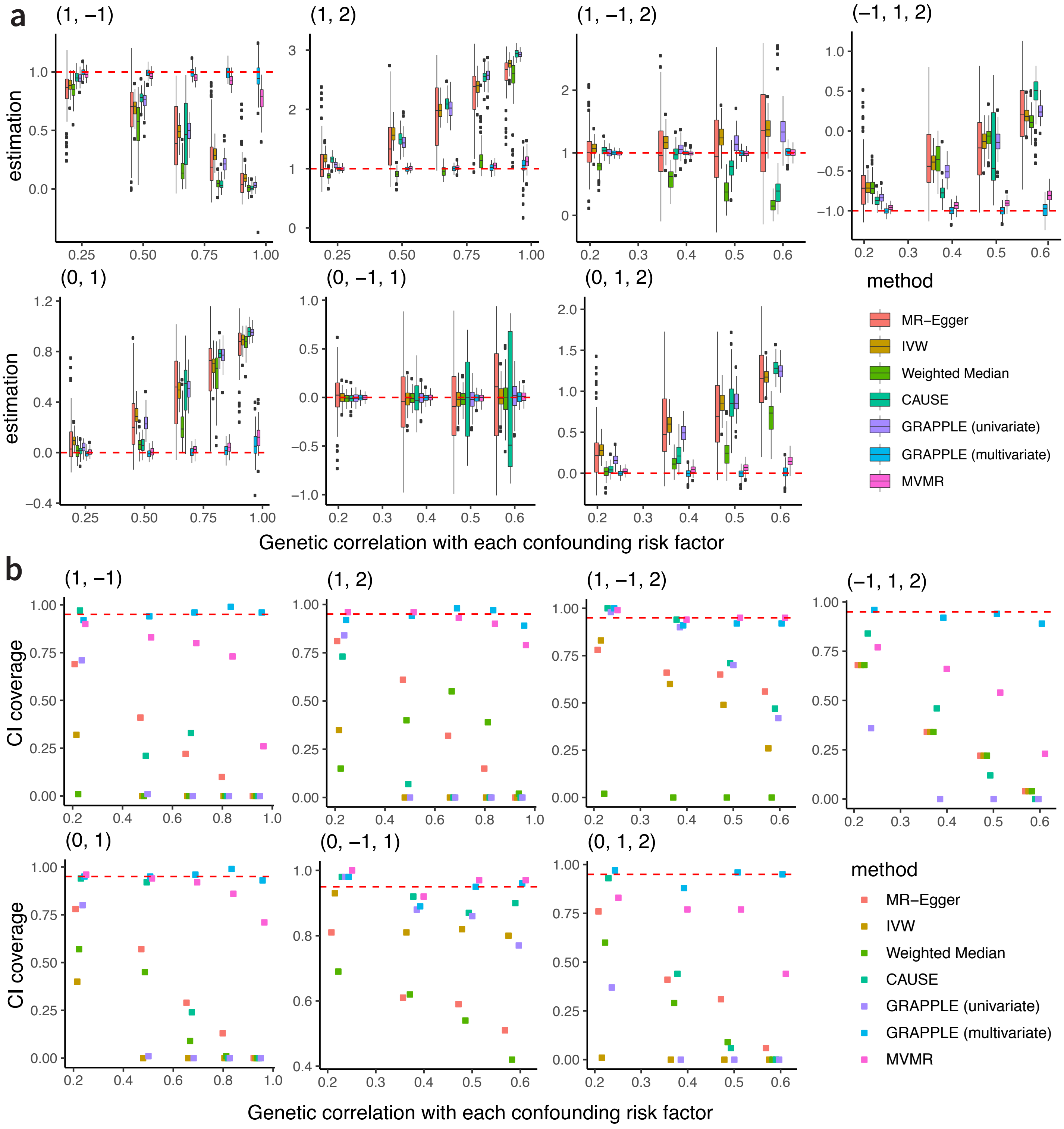


Figure S3: Comparison of different MR methods in settings with pleiotropic pathways when the selection threshold is $10^{-8}$ (top 106 SNPs). **a**, Boxplots of the estimated $\beta_1$ using different MR methods over 100 repeated random experiments. **b** The actual coverage of the 95% confidence intervals of $\beta_1$ provided by different methods. For CAUSE, we report the coverage of the 95% credible intervals of $\beta_1$. The red dotted line shows the expected 0.95 nominal level. For the three settings in the second row with $\beta_1 = 0$, the CI coverage is the same as $1 -$ type I error.

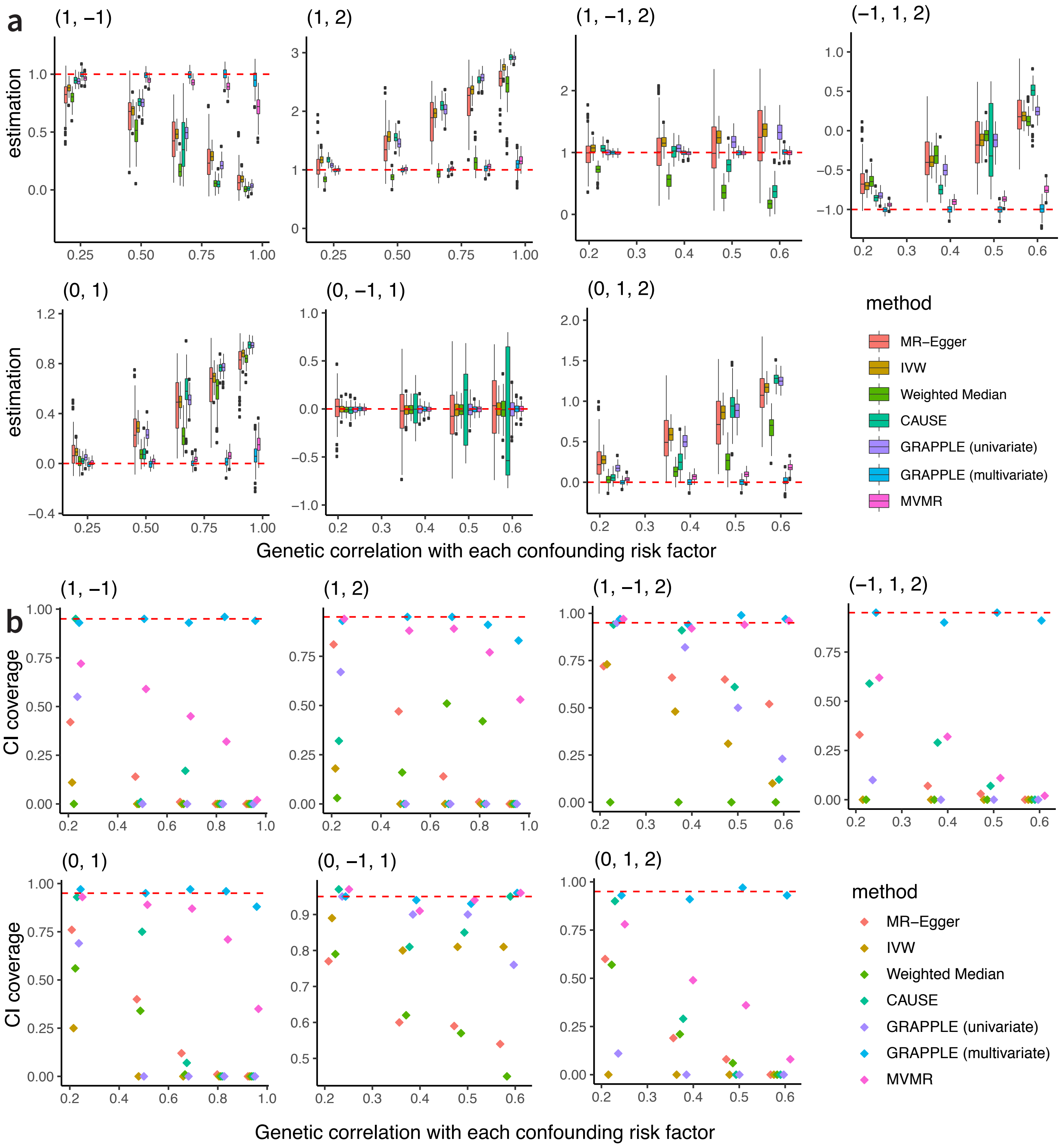


Figure S4: Same as Figure S3 with the selection threshold being $10^{-5}$ (top 217 SNPs).

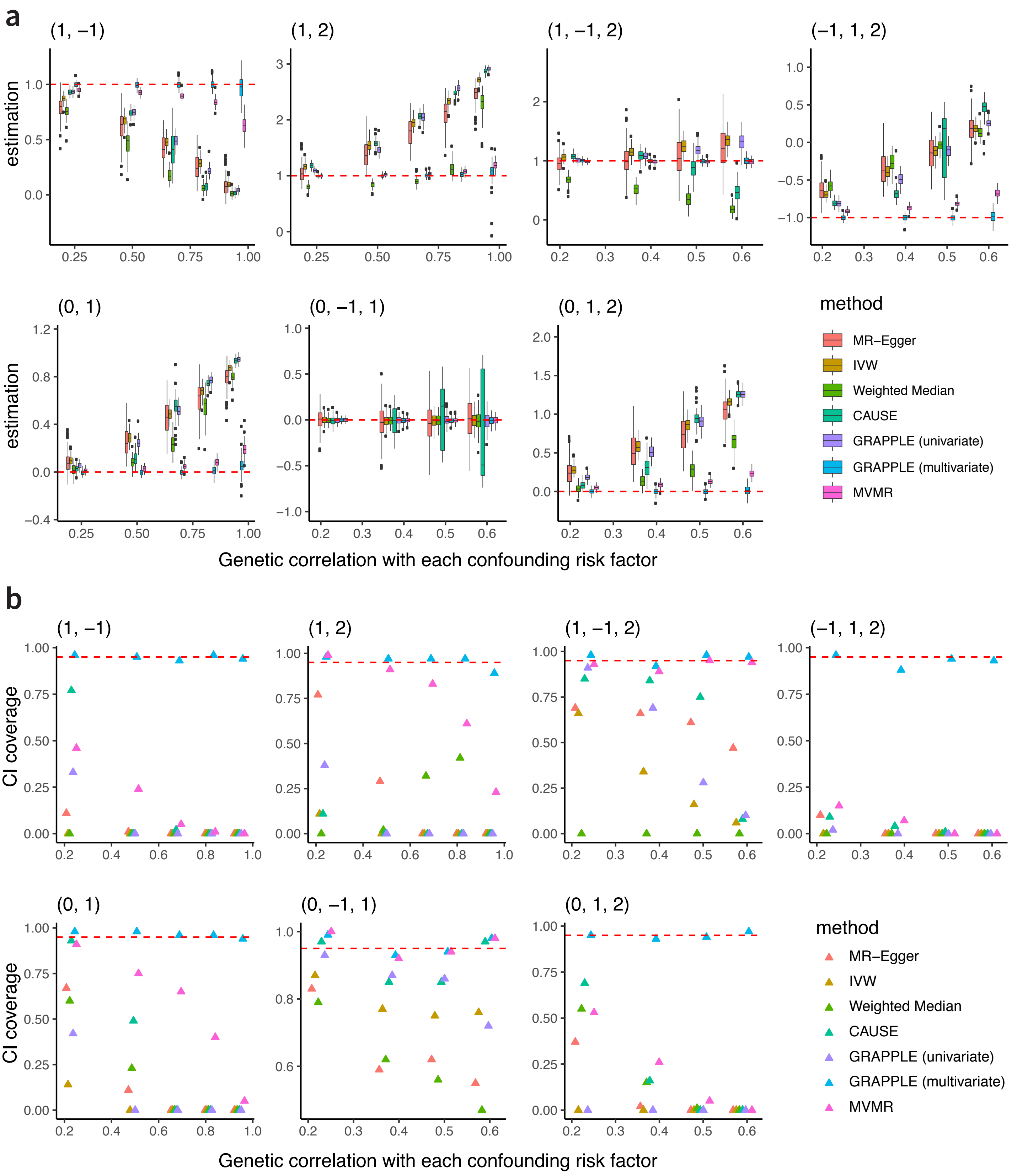


Figure S5: Same as Figure S3 with the selection threshold being 0.01 (top 422 SNPs).

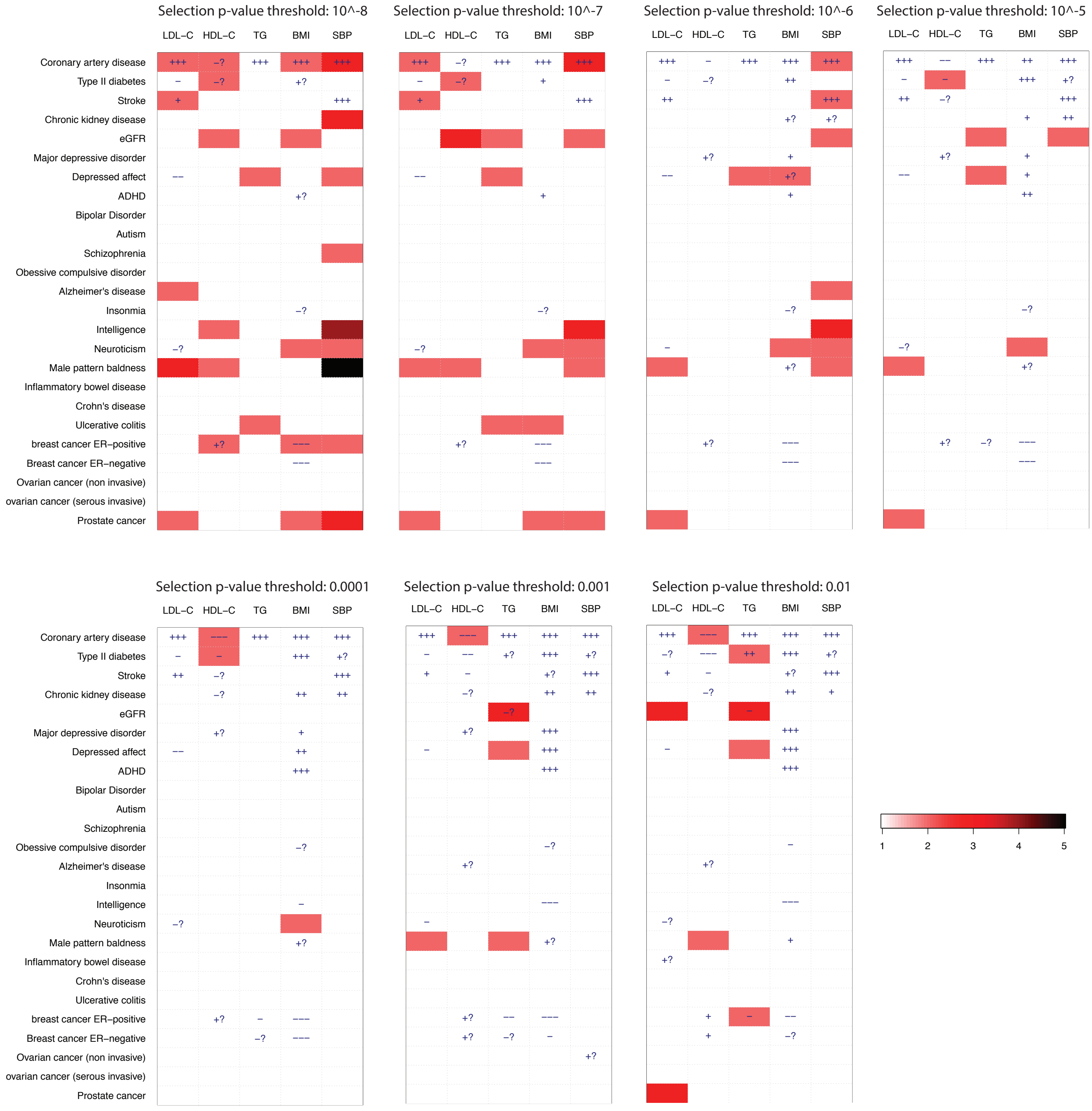


Figure S6: Landscape of pleiotropic pathways on 25 diseases. Each figure is for results obtained using one of the 7 p-value thresholds. The colors show the number of detected modes. The "+" sign shows a positive estimated effect and "−" sign shows a negative estimated effect.

### 2 Simulations

#### 2.1 The simulation model

Our simulation starts with the equation for each SNP $j$

$$\Gamma_j = \gamma_{j1}\beta_1 + \sum_{k=2}^{K} \gamma_{jk}\beta_k + \alpha_j$$

where we are interested in estimating $\beta_1$, the causal effect of our target risk factor $X_1$. We assume that $\alpha_j \overset{i.i.d.}{\sim} N(0, \tau^2)$ following the inSIDE assumption, while $\gamma_{jk}$ are correlated with $\gamma_{j1}$ for $k = 2, 3, \cdots, K$ due to the genetic correlations between $X_1$ and the confounding unmeasured risk factors $X_2, \cdots, X_K$. Specifically, we assume that for each $k = 2, 3, \cdots, K$,

$$\gamma_{jk} = \eta_{jk}\gamma_{j1} + (1 - \eta_{jk})\delta_{jk}$$

where

$$\eta_{jk} \overset{i.i.d.}{\sim} \text{Bernoulli}(\pi_k), \quad \delta_{jk} \overset{i.i.d.}{\sim} N(0, \tau_k^2)$$

and the data are the summary statistics $\hat{\Gamma}_j$ and $\hat{\gamma}_{1j}$ where

$$\hat{\Gamma}_j \overset{ind.}{\sim} N(\Gamma_j, \sigma_{Yj}^2), \quad \hat{\gamma}_{j1} \overset{ind.}{\sim} N(\gamma_{j1}, \sigma_{Xj}^2).$$

#### 2.2 The simulation settings

We base on real data estimations to set realistic values of $\gamma_{j1}$ as well as the standard errors $\sigma_{Y_j}$ and $\sigma_{X_j}$ in our simulations. Specifically, we take the BMI as the risk factor and SBP as the disease. With three-sample design, the BMI summary statistics from the GIANT consortium are used for SNP selection, and data from the UK Biobank for the two traits are used for estimation (see Table S1). With a p-value threshold of 0.01, we selected 786 independent SNPs. We treat the estimated $\widehat{\gamma}_{j1}$ for these SNPs as the true marginal associations $\gamma_{j1}$. The standard errors $\sigma_{Y_j}$ and $\sigma_{X_j}$ are the same as obtained from the GWAS summary statistics. We set $\tau^2 = \sum_j \gamma_{j1}^2/5$ for size of the uncorrelated pleiotropy and $\tau_2^2 = \cdots = \tau_K^2 = \tau^2/2$ for the SNPs effects on confounding risk factors when the SNPs are not on the shared pleiotropic pathway. Instead of using original selection p-values, we redefine the "selection p-value" as $2[1 - \Phi(|\gamma_{j1}|/\sigma_{Xj})]$ representing the signal strengths. Here $\Phi(\cdot)$ is the cumulative density function of the standard normal distribution.

Given the above settings, different $\pi_k$ corresponds to a different genetic correlation [5] between $X_1$ and $X_k$ following:

$$\rho_g(X_1, X_k) = \frac{\pi_k}{\sqrt{\pi_k + (1 - \pi_k)/10}}$$

We examine 10 combinations of $(\beta_1, \cdots, \beta_K)$ where $K = 1$, 2 or 3. When there are pleiotropic pathways, we consider both the scenario that the confounding risk factor has same or opposite sign as the effect of the target risk factor. Specifically, the ten combinations are:

- No pleiotropic pathways with $\beta_1 = 0.2$, 0.5 and 1.
- True causal effect is zero, one pleiotropic pathway: $(\beta_1, \beta_2) = (0, 1)$
- True causal effect is zero, with two pleiotropic pathways: $(\beta_1, \beta_2, \beta_3) = (0, -1, 1)$ or $(0, 1, 2)$;

- True causal effect is nonzero, one pleiotropic pathway: $(\beta_1, \beta_2) = (1, -1)$ or $(1, 2)$;
- True causal effect is nonzero, with two pleiotropic pathways: $(\beta_1, \beta_2, \beta_3) = (1, -1, 2)$ or $(-, 1, 2)$.

For each combination, we vary the values of $\pi_k$. When $K = 2$, there is only one pleiotropic pathway, and we set $\pi_2 = 0.1, 0.3, 0.5, 0.7, 0.9$. Here $\pi_2 = 0.1$ indicates that $\rho_g(X_1, X_2)$ is as small as 0.23 and $\pi_2 = 0.9$ is for the genetic correlation to be as large as 0.94. For $K = 3$ where there are two pleiotropic pathways, we require $\eta_{jk} = 1$ in at most one $k$ and we set $\pi_2 = \pi_3 = 0.1, 0.2, 0.3, 0.4$. This means that the two pleiotropic pathways have 0 genetic correlation, while they have the same genetic correlation with $X_1$, ranging from 0.23 ($\pi_2 = \pi_3 = 0.1$) to 0.59 ($\pi_2 = \pi_3 = 0.4$). For each $\boldsymbol{\beta}$ and $\boldsymbol{\pi}$ combination, we randomly repeat the experiments by $B = 100$ times.

#### 2.3 The simulation results

We compare GRAPPLE with 4 different MR methods that only take into account 1 risk factor: MR-Egger, IVW, Weighted Median and CAUSE, as well as a recently proposed MR method MVMR [26] for joint MR analysis of multiple risk factors. In the settings where there are pleiotropic pathways, we compares two methods in GRAPPLE. One is GRAPPLE using only the GWAS summary statistics for the target risk factor, where we use MR-RAPS to estimate the causal effect. The other is to perform a multivariate MR analysis including summary statistics from both the target risk factor and confounding risk factors.

First, when there is no pleiotropic pathway, all MR methods are able to provide a good estimation of the true causal effects (Fig. S2a). Similar to want we have observed in the benchmarking results based on real data (Fig. 2a), we observe that the common MR methods MR-Egger, IVW and Weighted Median can provide conservatively biased estimation even with a stringent selection threshold. GRAPPLE can keep providing an accurate and unbiased estimate of the causal effect regardless of the selection threshold, with the accuracy increasing with a more relaxed selection threshold. CAUSE may sometimes overestimate the true causal effect, as what we have already seen in Fig. 2a.

Next, we examine the power of GRAPPLE in detecting multiple modes and finding marker genes for each mode. We use three measurements. One is the detection rate, which is the chance of detecting more than one mode in the robustified profile likelihood. The other two measures are the marker precision and recall. We compare the marker SNPs that GRAPPLE returns with the set of SNPs that truly belong to the pleiotropic pathways. As shown in Fig. S2b, the detection of multiple modes performs the best when $\pi_k$ is neither too large nor too small, so that each pathway has enough SNPs among the selected IVs to contribute to a mode. When we vary the selection thresholds, we find that GRAPPLE is most sensitive to detect the modes with a stringent p-value threshold ($10^{-8}$) where only strongly associated SNPs are selected. On the other hand, including weakly associated SNPs will increase the marker recall rate, making it more informative to identify the hidden risk factors once the modes are detected.

Then, we compare GRAPPLE with other methods when there are pleiotropic pathways (Fig. S3-S5), using SNPs selected by different selection thresholds. We use two metrics for evaluation: the accuracy in the estimation of the true causal effect, and the coverage of the 95% confidence intervals from each MR method. For CAUSE, we report the 95% credible interval. When the true causal effect $\beta_1 = 0$, the CI coverage is equivalent to one minus the type-I error in claiming a non-zero causal effect. In terms of the estimation of the true causal effects, we observe that all univariate MR methods using only summary statistics of the target risk factor can have large bias when there are pleiotropic pathways, especially when the confounding unmeasured risk factors are highly or even moderately correlated with the target risk factor. In terms of the CI coverage, all univariate MR methods are generally not reliable when there are pleiotropic pathways and result in a larger type-I error than expected. Among all 5 univariate MR methods, CAUSE has the best coverage when the confounding risk factors has a small genetic correlation with the target risk factor, a scenario where the assumptions in CAUSE are likely to hold. Though our univariate GRAPPLE can sometimes provide a less biased estimation of $\beta_1$ compared with other univariate MR methods when there are pleiotropic pathways, our CIs are always too optimistic. Fortunately, GRAPPLE also has high detection rate of multi-modality in these scenarios with reasonably high marker precision, so that we are aware of the existence of pleiotropic pathways and have information to identify them.

Finally, we compare the multivariate GRAPPLE with MVMR, both of which use summary statistics from both the target and confounding risk factors (Fig. S3-S5). Compared to the univariate MR methods, both methods are much more accurate in estimation and CI coverage. MVMR can suffer from weak instrument bias and is also sensitive to the genetic correlation between confounding risk factors and the target risk factor. In contrast, our multivariate GRAPPLE can keep providing accurate estimation of $\beta_1$, as well as reliable CIs with low type-I error regardless of either the genetic correlations of confounding risk factors or the SNPs selection thresholds.

### 3 Additional mathematical details

#### 3.1 Correlations of GWAS summary statistics from overlapping cohorts

We provide detailed calculations for the derivation of Equation (8). For any shared sample $s$, let

$$Y_s = \Gamma_j Z_{js} + \epsilon_{js}, \quad X_{ks} = \gamma_{jk} Z_{js} + e_{jks}$$

where $Y_s$, $X_{ks}$ and $Z_{js}$ all have mean 0 and the $Z_{js}$ has variance 1 for convenience. Then as for most SNPs, its individual genetic effect is very small, with $\Gamma_j = o(1)$ and $\gamma_{jk} = o(1)$ we have

$$\mathrm{Cov}\left[Y_s, X_{ks}\right] = \Gamma_j \gamma_{jks} \mathrm{Var}\left[Z_{js}\right] + \mathrm{Cov}\left[\epsilon_{js}, e_{jks}\right] \approx \mathrm{Cov}\left[\epsilon_{js}, e_{jks}\right]$$

The summary statistics are computed from marginal regression, so we have

$$\widehat{\gamma}_j = \frac{\widehat{\mathrm{Cov}}_{n_X}(X, Z_j)}{\widehat{\mathrm{Cov}}_{n_X}(Z_j, Z_j)} = \widehat{\mathrm{Cov}}_{n_X}(X, Z_j), \ \widehat{\Gamma}_j = \frac{\widehat{\mathrm{Cov}}_{n_Y}(Y, Z_j)}{\widehat{\mathrm{Cov}}_{n_Y}(Z_j, Z_j)} = \widehat{\mathrm{Cov}}_{n_Y}(Y, Z_j).$$

where $\widehat{\mathrm{Cov}}_n$ is the sample covariance operator.

Thus,

$$\begin{aligned}
\mathrm{Cov}\left[\widehat{\Gamma}_j, \widehat{\gamma}_{jk}\right] &= \frac{1}{N_{ek} N_o} \mathrm{Cov}\left[\sum_{s=1}^{N_{sk}} Y_s Z_{js}, \sum_{s=1}^{N_{sk}} X_{sk} Z_{js}\right] \\
&= \frac{N_{sk}}{N_{ek} N_o} \mathrm{Cov}\left[Y_s Z_{js}, X_{sk} Z_{js}\right] \\
&= \frac{N_{sk}}{N_{ek} N_o} \left(\Gamma_j \gamma_{jk} \mathrm{Var}\left[Z_{js}^2\right] + \mathrm{Var}\left[Z_{js}\right] \mathrm{Cov}\left[\epsilon_{js}, e_{jks}\right]\right) \\
&\approx \frac{N_{sk}}{N_{ek} N_o} \mathrm{Var}\left[Z_{js}\right] \mathrm{Cov}\left[\epsilon_{js}, e_{jks}\right]
\end{aligned}$$

and

$$\begin{aligned}
\mathrm{Var}\left[\widehat{\Gamma}_j\right] &= \frac{1}{N_o} \mathrm{Var}\left[Y_s Z_{js}\right] \approx \frac{1}{N_o} \mathrm{Var}\left[Z_{js}\right] \mathrm{Var}\left[\epsilon_{js}\right] \\
\mathrm{Var}\left[\widehat{\gamma}_{jk}\right] &= \frac{1}{N_{ek}} \mathrm{Var}\left[Y_s Z_{js}\right] \approx \frac{1}{N_{ek}} \mathrm{Var}\left[Z_{js}\right] \mathrm{Var}\left[e_{jks}\right] \\
\mathrm{Var}\left[Y_s\right] &\approx \mathrm{Var}\left[\epsilon_{js}\right], \quad \mathrm{Var}\left[X_{ks}\right] \approx \mathrm{Var}\left[e_{jks}\right]
\end{aligned}$$

Above approximations show that

$$\mathrm{Corr}\left[\widehat{\Gamma}_j, \widehat{\gamma}_{jk}\right] \approx \frac{N_{sk}}{\sqrt{N_{ek} N_o}} \mathrm{Corr}\left[Y_s, X_{ks}\right]$$

#### 3.2 Robustified profile likelihood methods for multivariate MR

With the random effect model on $\alpha_j$, we further have

$$\begin{pmatrix} \widehat{\Gamma}_j \\ \widehat{\boldsymbol{\gamma}}_j \end{pmatrix} \sim \mathcal{N}\left(\begin{pmatrix} \boldsymbol{\gamma}_j^T \boldsymbol{\beta} \\ \boldsymbol{\gamma}_j \end{pmatrix}, \mathrm{diag}\begin{pmatrix} \sigma_{Y_j} \\ \sigma_{X_j} \end{pmatrix} \Sigma \ \mathrm{diag}\begin{pmatrix} \sigma_{Y_j} \\ \sigma_{X_j} \end{pmatrix} + \begin{pmatrix} \tau^2 & \mathbf{0} \\ \mathbf{0} & \mathbf{0} \end{pmatrix}\right)$$

Then, we estimate the parameters $\boldsymbol{\beta}$ and $\tau^2$ using $K+1$ estimation equations. Recall that we have defined

$$t_j(\boldsymbol{\beta}, \tau^2) = \frac{\widehat{\Gamma}_j - \widehat{\boldsymbol{\gamma}}_j{}^T \boldsymbol{\beta}}{\sqrt{\sigma_{Y_j}^2 + \boldsymbol{\beta}^T \Sigma_{\boldsymbol{X}_j} \boldsymbol{\beta} - 2\boldsymbol{\beta}^T \Sigma_{\boldsymbol{X}_j Y_j} + \tau^2}} \tag{1}$$

and the robust profile likelihood is the optimization function

$$l(\boldsymbol{\beta}, \tau^2) = -\sum_j l_j(\boldsymbol{\beta}, \tau^2) = -\sum_j \rho\left(t_j(\boldsymbol{\beta}, \tau^2)\right) \tag{2}$$

The estimation equations for $\boldsymbol{\beta}$ are from the derivatives of the robust profile likelihood:

$$\boldsymbol{\varphi}_1(\boldsymbol{\beta}, \tau^2) = \frac{\partial l(\boldsymbol{\beta}, \tau^2)}{\partial \boldsymbol{\beta}} = \mathbf{0}$$

The estimation equation for $\tau^2$ is

$$\varphi_2(\boldsymbol{\beta}, \tau^2) = l(\boldsymbol{\beta}, \tau^2) - p\eta = 0$$

where $\eta = \mathbb{E}\left[\rho(Z)\right]$ with $Z \sim \mathcal{N}(0,1)$. Let $\boldsymbol{\varphi}(\cdot) = \begin{pmatrix} \boldsymbol{\varphi}_1(\cdot) \\ \varphi_2(\cdot) \end{pmatrix}$.

We use delta method to calculate the asymptotic distribution of $\widehat{\boldsymbol{\beta}}$ and $\widehat{\tau}^2$ obtained from the above estimation equations. To make it clear, we denote the true values of $\boldsymbol{\beta}$ and $\tau$ as $\boldsymbol{\beta}_0$ and $\tau_0$. Then, we have

$$\mathbf{0} = \boldsymbol{\varphi}(\widehat{\boldsymbol{\beta}}, \widehat{\tau}^2) \approx \boldsymbol{\varphi}(\boldsymbol{\beta}_0, \tau_0^2) + \dot{\boldsymbol{\varphi}}(\boldsymbol{\beta}_0, \tau_0^2) \begin{pmatrix} \widehat{\boldsymbol{\beta}} - \boldsymbol{\beta}_0 \\ \widehat{\tau}_0^2 - \tau_0^2 \end{pmatrix}$$

Let

$$A = \mathbb{E}\left[-\dot{\boldsymbol{\varphi}}(\boldsymbol{\beta}_0, \tau_0^2)\right], \; B = \mathrm{Var}\left[\boldsymbol{\varphi}(\boldsymbol{\beta}_0, \tau_0^2)\right]$$

Then if $\widehat{\boldsymbol{\beta}}$ and $\widehat{\tau}^2$ are consistent estimates, we would have asymptotically

$$\begin{pmatrix} \widehat{\boldsymbol{\beta}} - \boldsymbol{\beta}_0 \\ \widehat{\tau}_0^2 - \tau_0^2 \end{pmatrix} \sim \mathcal{N}(0, A^{-1}BA^{-T})$$

Thus, we only need to estimate $A$ and $B$.

Let's first discuss how to estimate $B$. Notice that

$$\boldsymbol{\varphi}_{1j}(\boldsymbol{\beta}, \tau^2) = \rho'(t_j) \frac{\partial t_j}{\partial \boldsymbol{\beta}}.$$

Based on the definition of $t_j$ in (1), we have

$$\frac{\partial t_j}{\partial \boldsymbol{\beta}} = -\frac{(\sigma_{Y_j}^2 + \beta^T \Sigma_{X_j} \boldsymbol{\beta} - 2\boldsymbol{\beta}^T \Sigma_{X_j Y_j} + \tau^2)\widehat{\boldsymbol{\gamma}}_j + (\widehat{\Gamma}_j - \widehat{\boldsymbol{\gamma}}_j{}^T \boldsymbol{\beta})(\Sigma_{X_j} \boldsymbol{\beta} - \Sigma_{X_j Y_j})}{\left(\sigma_{Y_j}^2 + \boldsymbol{\beta}^T \Sigma_{X_j} \boldsymbol{\beta} - 2\boldsymbol{\beta}^T \Sigma_{X_j Y_j} + \tau^2\right)^{3/2}}$$

As

$$\mathrm{Var}\left(\widehat{\Gamma}_j - \widehat{\boldsymbol{\gamma}}_j{}^T \boldsymbol{\beta}_0\right) = \sigma_{Y_j}^2 + \boldsymbol{\beta}_0{}^T \Sigma_{X_j} \boldsymbol{\beta}_0 - 2\boldsymbol{\beta}^T \Sigma_{X_j Y_j} + \tau_0^2$$

and

$$\mathrm{Cov}\left(\widehat{\boldsymbol{\gamma}}_j, \widehat{\Gamma}_j - \widehat{\boldsymbol{\gamma}}_j{}^T \boldsymbol{\beta}_0\right) = \Sigma_{X_j Y_j} - \Sigma_{X_j} \boldsymbol{\beta}_0$$

We have,

$$\text{Cov}\left(t_j, \frac{\partial t_j}{\partial \boldsymbol{\beta}}\right) = 0$$

at the true values $\boldsymbol{\beta}_0$ and $\tau_0$. Further because $t_j$ and $\frac{\partial t_j}{\partial \boldsymbol{\beta}}$ are linear transformations of $\widehat{\Gamma}_j$ and $\widehat{\boldsymbol{\gamma}}_j$, they are jointly Gaussian. Thus, $t_j$ and $\frac{\partial t_j}{\partial \boldsymbol{\beta}}$ are independent at the true values $\boldsymbol{\beta}_0$ and $\tau_0$. So we have

$$\begin{aligned}
\text{Var}\left[\boldsymbol{\varphi}_1(\boldsymbol{\beta}_0, \tau_0^2)\right] &= \sum_j \text{Var}\left[\boldsymbol{\varphi}_{1j}(\boldsymbol{\beta}_0, \tau_0^2)\right] \\
&= \sum_j \mathbb{E}\left[\boldsymbol{\varphi}_{1j}(\boldsymbol{\beta}_0, \tau_0^2)^2\right] \\
&= \sum_j \mathbb{E}\left[\rho'(t_j(\boldsymbol{\beta}_0, \tau_0^2))^2\right] \mathbb{E}\left[\frac{\partial t_j}{\partial \boldsymbol{\beta}}(\boldsymbol{\beta}_0, \tau_0^2)\frac{\partial t_j}{\partial \boldsymbol{\beta}}(\boldsymbol{\beta}_0, \tau_0^2)^T\right] \\
&= \mathbb{E}\left[\rho'(Z)^2\right] \sum_j \mathbb{E}\left[\frac{\partial t_j}{\partial \boldsymbol{\beta}}(\boldsymbol{\beta}_0, \tau_0^2)\frac{\partial t_j}{\partial \boldsymbol{\beta}}(\boldsymbol{\beta}_0, \tau_0^2)^T\right]
\end{aligned}$$

Also, we have

$$\text{Cov}\left[\boldsymbol{\varphi}_{1j}(\boldsymbol{\beta}_0, \tau_0^2), \varphi_{2j}(\boldsymbol{\beta}_0, \tau_0^2)\right] = \sum_j \mathbb{E}\left[\rho'(t_j(\boldsymbol{\beta}_0, \tau_0^2))\frac{\partial t_j}{\partial \boldsymbol{\beta}}(\boldsymbol{\beta}_0, \tau_0^2)\rho'(t_j(\boldsymbol{\beta}_0, \tau_0^2))\right] = 0$$

and

$$\text{Var}\left[\varphi_{2j}(\boldsymbol{\beta}_0, \tau_0^2)\right] = p\,\text{Var}\left[\rho(Z)\right]$$

where $Z \sim N(0, 1)$. Thus, we can estimate $B$ as

$$\widehat{B} = \begin{pmatrix} \text{Var}\left[\rho(Z)\right] \sum_j \frac{\partial t_j}{\partial \boldsymbol{\beta}}(\widehat{\boldsymbol{\beta}}, \widehat{\tau}^2)\frac{\partial t_j}{\partial \boldsymbol{\beta}}(\widehat{\boldsymbol{\beta}}, \widehat{\tau}^2)^T & 0 \\ 0 & p\,\text{Var}\left[\rho(Z)\right] \end{pmatrix}$$

Next, we estimate $A$. We need to calculate the function of $\dot{\boldsymbol{\varphi}}(\boldsymbol{\beta}, \tau^2)$. we have

$$\begin{aligned}
\frac{\partial \boldsymbol{\varphi}_{1j}}{\partial \boldsymbol{\beta}} &= \rho''(t_j)\left(\frac{\partial t_j}{\partial \boldsymbol{\beta}}\right)\left(\frac{\partial t_j}{\partial \boldsymbol{\beta}}\right)^T + \rho'(t_j)\frac{\partial^2 t_j}{\partial \boldsymbol{\beta}\partial \boldsymbol{\beta}} \\
\frac{\partial \boldsymbol{\varphi}_{1j}}{\partial \tau^2} &= \rho''(t_j)\left(\frac{\partial t_j}{\partial \tau^2}\right)\left(\frac{\partial t_j}{\partial \boldsymbol{\beta}}\right) + \rho'(t_j)\frac{\partial^2 t_j}{\partial \tau^2 \partial \boldsymbol{\beta}} \\
\frac{\partial \varphi_{2j}}{\partial \boldsymbol{\beta}} &= \boldsymbol{\varphi}_{1j} \\
\frac{\partial \varphi_{2j}}{\partial \tau^2} &= \rho'(t_j)\frac{\partial t_j}{\partial \tau^2}
\end{aligned}$$

As

$$\frac{\partial t_j}{\partial \tau^2} = -\frac{1}{2}\frac{t_j}{\sigma_{Y_j}^2 + \boldsymbol{\beta}^T \Sigma_{X_j} \boldsymbol{\beta} - 2\boldsymbol{\beta}^T \Sigma_{X_j Y_j} + \tau^2}$$

we have

$$\mathbb{E}\left[\frac{\partial \varphi_{2j}}{\partial \tau^2}(\boldsymbol{\beta}_0, \tau_0^2)\right] = -\frac{\mathbb{E}\left[\rho'(Z)Z\right]}{2} \sum_j \frac{1}{\sigma_{Y_j}^2 + \boldsymbol{\beta}_0^T \Sigma_{X_j} \boldsymbol{\beta}_0 - 2\boldsymbol{\beta}_0^T \Sigma_{X_j Y_j} + \tau_0^2}$$

$$\begin{aligned}\mathbb{E}\left[\frac{\partial \boldsymbol{\varphi}_{1j}}{\partial \tau^2}(\boldsymbol{\beta}_0, \tau_0^2)\right] &= \mathbb{E}\left[\rho'(t_j(\boldsymbol{\beta}_0, \tau_0^2))\frac{\partial^2 t_j}{\partial \tau^2 \partial \boldsymbol{\beta}}(\boldsymbol{\beta}_0, \tau_0^2)\right] \\ &= \frac{\mathbb{E}\left[\rho'(Z)Z\right]}{2}\sum_j \frac{\Sigma_{X_j}\boldsymbol{\beta} - \Sigma_{X_jY_j}}{(\sigma_{Y_j}^2 + \boldsymbol{\beta}_0^T\Sigma_{X_j}\boldsymbol{\beta}_0 - 2\boldsymbol{\beta}_0^T\Sigma_{X_jY_j} + \tau_0^2)^2}\end{aligned}$$

We also have

$$\mathbb{E}\left[\frac{\partial \varphi_{2j}}{\partial \boldsymbol{\beta}}(\boldsymbol{\beta}_0, \tau_0^2)\right] = \mathbf{0}$$

$$\begin{aligned}\mathbb{E}\left[\frac{\partial \boldsymbol{\varphi}_{1j}}{\partial \boldsymbol{\beta}}(\boldsymbol{\beta}_0, \tau_0^2)\right] =& \mathbb{E}\left[\rho''(Z)\right]\mathbb{E}\left[\sum_j \frac{\partial t_j}{\partial \boldsymbol{\beta}}(\boldsymbol{\beta}_0, \tau_0^2)\frac{\partial t_j}{\partial \boldsymbol{\beta}}(\boldsymbol{\beta}_0, \tau_0^2)^T\right] + \\ & \mathbb{E}\left[\sum_j \rho'(t_j(\boldsymbol{\beta}_0, \tau_0^2))\frac{\partial^2 t_j}{\partial \boldsymbol{\beta}\partial \boldsymbol{\beta}}(\boldsymbol{\beta}_0, \tau_0^2)\right]\end{aligned}$$

where we can estimate the last two expectations on the right hand side by the corresponding sample means.

### 4 Data resources

Below, we list the datasets that we have used in the paper and where their download sources. Table S1 and table S2 summarize how these datasets are used in different analyses in the paper.

| Analysis | Risk factor datasets | | Disease dataset |
|---|---|---|---|
| | Selection | Estimation | |
| Validation1: | | | |
| BMI → BMI | BMI-ukb | BMI-giant17eu-F | BMI-giant17eu-M |
| T2D → T2D | T2D-ukb | T2D-diagram12-F | T2D-diagram12-M |
| Height → Height | Height-ukb | Height-giant13-F | Height-giant13-M |
| Validation2: | | | |
| BMI → T2D | BMI-ukb | BMI-giant17eu | T2D-diagram12-Im |
| LDL-C → CAD | LDL-gera18 | LDL-glgc13 | CAD-Nelson17 |
| Height → Smoking | Height-giant14 | Height-ukb | Smoking-ukb19 |
| SBP → Stroke | SBP-gera17 | SBP-ukb | AS-Malik18eu |
| Validation3: | | | |
| BMI → T2D | BMI-ukb | BMI-giant17eu-F/M | T2D-diagram-F/M |
| T2D → BMI | T2D-ukb | T2D-diagram12-F/M | BMI-giant17eu-F/M |
| LDL-C → CAD | LDL-gera18 | LDL-glgc13 | CAD-Nelson17 |
| CAD → LDL-C | CAD-c4d11 | CAD-CARDIoGRAM11 | LDL-glgc13 |
| Validation4: | | | |
| CRP → CAD | CRP-Prins17 | CRP-Dehghan11 | CAD-Nelson17 |
| CRP + LDL-C → CAD | (CRP-Prins17, LDL-gera18) | (CRP-Dehghan11, LDL-glgc13) | CAD-Nelson17 |
| Screening risk factors: | | | |
| LDL-C | LDL-gera18 | LDL-glgc13 | - |
| HDL-C | HDL-gera18 | HDL-glgc13 | - |
| TG-C | TG-gera18 | TG-glgc13 | - |
| BMI | BMI-JapB | BMI-ukb | - |
| SBP | SBP-gera17 | SBP-ukb | - |
| Simulation: | | | |
| BMI → SBP | BMI-giant17eu | BMI-ukb | SBP-ukb |

Table S1: Names of the datasets used in validation analyses and screening risk factors

- BMI-ukb: downloaded from UK Biobank Neale's lab [12] with phenotype code 21001.
- BMI-giant17eu-F, BMI-giant17eu-M, BMI-giant17eu: downloaded from GIANT consortium website [2], 2017 adjusted for smoking data.

| Screening Disease | dataset | Screening Disease | dataset |
|---|---|---|---|
| CAD | CAD-Nelson17 | Insomnia | Insomnia-ukb19 |
| Type 2 diabetes | T2D-diagram12-Im | Intelligence | IQ-Savage18 |
| Stroke | AS-Malik18eu | Neuroticism | Neuro-Hill19 |
| Chronic kidney disease | CKD-Wuttke19 | Male pattern baldness | MBP-ukb19 |
| eGFR | eGFR-Wuttke19 | IBD | IBD-Liu15 |
| MDD | MDD-PGC18 | Crohn's disease | CR-Liu15 |
| Depressed affect | Dep-Nagel18 | Ulcerative colitis | UC-Liu15 |
| ADHD | ADHD-pgc19 | Breast cancer ER+ | Breast-Micha7erp |
| Bipolar Disorder | BIP-pgc19 | Breast cancer ER- | Breast-Micha17ern |
| Autism | Autism-pgc17 | Ovarian cancer (non-invasive) | Ovarian-Phelan17ni |
| Schizophrenia | SCZ-pgc13 | | |
| Obsessive compulsive disorder | OCD-pgc18 | Ovarian cancer (serous invasive) | Ovarian-Phelan17si |
| Alzheimer | Alzhe-Marioni18 | Prostate cancer | Prostate-ellipse18 |

Table S2: Names of the datasets used for the 25 disesases in the screening application

- BMI-JapB: data originally from [3] and downloaded from GWAS Catalog with study accession number GCST004904.
- T2D-diagram12-F, T2D-diagram12-M, T2D-diagram12-Im: data originally from [19] and downloaded from `https://diagram-consortium.org/downloads.html`.
- T2D-ukb: downloaded from UK Biobank Neale's lab [12] with phenotype code 20002-1223.
- Height-giant13-F, Height-giant13-M: data originally from [24] and downloaded from GIANT consortium website [2].
- Height-giant14: data originally from [32] and downloaded from GIANT consortium website [2].
- Height-ukb: downloaded from UK Biobank Neale's lab [12] with phenotype code 50.
- LDL-glgc13, HDL-glgc13, TG-glgc13: data originally from [31] and downloaded from `http://csg.sph.umich.edu/willer/public/lipids2013/` (Joint analysis of metoboship and GWAS data).
- LDL-gera18, HDL-gera18, TG-gera18: data originally from [11] and downloaded from GWAS Catalog with study accession numbers GCST007141, GCST007140 and GCST007142.
- SBP-gera17: data originally from [10] and downloaded from GWAS Catalog with study accession number GCST007095.
- SBP-ukb: downloaded from UK Biobank Neale's lab [12] with phenotype code 4080.
- CAD-Nelson17: data originally from [21] and downloaded from GWAS Catalog with study accession number GCST004787.
- CAD-c4d11: data originally from [6] and downloaded from `http://www.cardiogramplusc4d.org/data-downloads/`

- CAD-CARDIoGRAM11: data originally from [29] and downloaded from `http://www.cardiogramplusc4d.org/data-downloads/`
- CRP-Prins17: data originally from [23] and downloaded from GWAS Catalog with study accession number GCST005067.
- CRP-Dehghan11: Data from [7] and requested from original authors.
- AS-Malik18EU: data originally from [16] and downloaded from GWAS Catalog with study accession number GCST006906.
- Smoking-ukb19: data originally from [14] and downloaded from GWAS Catalog with study accession number GCST007327.
- CKD-Wuttke19, eGFR-Wuttke19: data originally from [34] and downloaded from GWAS Catalog with study accession numbers GCST008065 and GCST008059.
- Dep-Nagel18: data originally from [20] and downloaded from GWAS Catalog with study accession number GCST006475.
- ADHD-pgc19: data originally from [8] and downloaded from the PGC website `https://www.med.unc.edu/pgc/download-results/`.
- MDD-pgc18: data originally from [33] and downloaded from the PGC website `https://www.med.unc.edu/pgc/download-results/`.
- BIP-pgc19: data originally from [30] and downloaded from the PGC website `https://www.med.unc.edu/pgc/download-results/`.
- Autism-pgc17: data originally from [1] and downloaded from the PGC website `https://www.med.unc.edu/pgc/download-results/`.
- SCZ-pgc13: data originally from [25] and downloaded from the PGC website `https://www.med.unc.edu/pgc/download-results/`.
- OCD-pgc18: data originally from [4] and downloaded from the PGC website `https://www.med.unc.edu/pgc/download-results/`.
- Alzhe-Marioni18: data originally from [17] and downloaded from GWAS Catalog with study accession number GCST005922.
- Neuro-Hill19: data originally from [9] and downloaded from GWAS Catalog with study accession number GCST007710.
- IQ-Savage18: data originally from [27] and downloaded from GWAS Catalog with study accession number GCST006250.
- Insomnia-ukb19: data originally from [13] and downloaded from GWAS Catalog with study accession number GCST007387.
- MBP-ukb19: data originally from [35] and downloaded from GWAS Catalog with study accession number GCST007020.

- IBD-Liu15, UC-Liu15, CR-Liu15: data originally from [15] and downloaded from GWAS Catalog with study accession numbers GCST003043, GCST003045 and GCST003044.

- Breast-Micha17erp, Breast-Micha17ern: data originally from [18] and downloaded from GWAS Catalog with study accession number GCST004988.

- Ovarian-Phelan17ni, Ovarian-Phelan17si: data originally from [22] and downloaded from GWAS Catalog with study accession numbers GCST004462 and GCST004478.

- Prostate-ellipse18: data originally from [28] and downloaded from GWAS Catalog with study accession number GCST006085

### References


[1] Meta-analysis of gwas of over 16,000 individuals with autism spectrum disorder highlights a novel locus at 10q24. 32 and a significant overlap with schizophrenia. *Molecular autism*, 8:1–17, 2017.

[2] *GIANT consortium data files*, (accessed 2020/3/25). `https://portals.broadinstitute.org/collaboration/giant/index.php/GIANT_consortium_data_files`.

[3] M. Akiyama, Y. Okada, M. Kanai, A. Takahashi, Y. Momozawa, M. Ikeda, N. Iwata, S. Ikegawa, M. Hirata, K. Matsuda, et al. Genome-wide association study identifies 112 new loci for body mass index in the japanese population. *Nature genetics*, 49(10):1458, 2017.

[4] P. D. Arnold, K. D. Askland, C. Barlassina, L. Bellodi, O. Bienvenu, D. Black, M. Bloch, H. Brentani, C. L. Burton, B. Camarena, et al. Revealing the complex genetic architecture of obsessive-compulsive disorder using meta-analysis. *Molecular psychiatry*, 23(5):1181–1181, 2018.

[5] B. Bulik-Sullivan, H. K. Finucane, V. Anttila, A. Gusev, F. R. Day, P.-R. Loh, L. Duncan, J. R. Perry, N. Patterson, E. B. Robinson, et al. An atlas of genetic correlations across human diseases and traits. *Nature genetics*, 47(11):1236, 2015.

[6] Coronary Artery Disease (C4D) Genetics Consortium et al. A genome-wide association study in europeans and south asians identifies five new loci for coronary artery disease. *Nature genetics*, 43(4):339, 2011.

[7] A. Dehghan, J. Dupuis, M. Barbalic, J. C. Bis, G. Eiriksdottir, C. Lu, N. Pellikka, H. Wallaschofski, J. Kettunen, P. Henneman, et al. Meta-analysis of genome-wide association studies in¿ 80 000 subjects identifies multiple loci for c-reactive protein levelsclinical perspective. *Circulation*, 123(7):731–738, 2011.

[8] D. Demontis, R. K. Walters, J. Martin, M. Mattheisen, T. D. Als, E. Agerbo, G. Baldursson, R. Belliveau, J. Bybjerg-Grauholm, M. Bækvad-Hansen, et al. Discovery of the first genome-wide significant risk loci for attention deficit/hyperactivity disorder. *Nature genetics*, 51(1):63–75, 2019.

[9] W. D. Hill, A. Weiss, D. C. Liewald, G. Davies, D. J. Porteous, C. Hayward, A. M. McIntosh, C. R. Gale, and I. J. Deary. Genetic contributions to two special factors of neuroticism are associated with affluence, higher intelligence, better health, and longer life. *Molecular psychiatry*, pages 1–19, 2019.

[10] T. J. Hoffmann, G. B. Ehret, P. Nandakumar, D. Ranatunga, C. Schaefer, P.-Y. Kwok, C. Iribarren, A. Chakravarti, and N. Risch. Genome-wide association analyses using electronic health records identify new loci influencing blood pressure variation. *Nature genetics*, 49(1):54, 2017.

[11] T. J. Hoffmann, E. Theusch, T. Haldar, D. K. Ranatunga, E. Jorgenson, M. W. Medina, M. N. Kvale, P.-Y. Kwok, C. Schaefer, R. M. Krauss, et al. A large electronic-health-record-based genome-wide study of serum lipids. *Nature genetics*, 50(3):401–413, 2018.

[12] N. lab. *UK Biobank round 1 GWAS results*, 2017 (accessed 2020/3/25). `http://www.nealelab.is/blog/2017/7/19/rapid-gwas-of-thousands-of-phenotypes-for-337000-samples-in-the-uk-biobank`.

[13] J. M. Lane, S. E. Jones, H. S. Dashti, A. R. Wood, K. G. Aragam, V. T. van Hees, L. B. Strand, B. S. Winsvold, H. Wang, J. Bowden, et al. Biological and clinical insights from genetics of insomnia symptoms. *Nature genetics*, 51(3):387–393, 2019.

[14] R. K. Linnér, P. Biroli, E. Kong, S. F. W. Meddens, R. Wedow, M. A. Fontana, M. Lebreton, S. P. Tino, A. Abdellaoui, A. R. Hammerschlag, et al. Genome-wide association analyses of risk tolerance and risky behaviors in over 1 million individuals identify hundreds of loci and shared genetic influences. *Nature genetics*, 51(2):245–257, 2019.

[15] J. Z. Liu, S. Van Sommeren, H. Huang, S. C. Ng, R. Alberts, A. Takahashi, S. Ripke, J. C. Lee, L. Jostins, T. Shah, et al. Association analyses identify 38 susceptibility loci for inflammatory bowel disease and highlight shared genetic risk across populations. *Nature genetics*, 47(9):979, 2015.

[16] R. Malik, G. Chauhan, M. Traylor, M. Sargurupremraj, Y. Okada, A. Mishra, L. Rutten-Jacobs, A.-K. Giese, S. W. Van Der Laan, S. Gretarsdottir, et al. Multiancestry genome-wide association study of 520,000 subjects identifies 32 loci associated with stroke and stroke subtypes. *Nature genetics*, 50(4):524–537, 2018.

[17] R. E. Marioni, S. E. Harris, Q. Zhang, A. F. McRae, S. P. Hagenaars, W. D. Hill, G. Davies, C. W. Ritchie, C. R. Gale, J. M. Starr, et al. Gwas on family history of alzheimer's disease. *Translational psychiatry*, 8(1):1–7, 2018.

[18] K. Michailidou, S. Lindström, J. Dennis, J. Beesley, S. Hui, S. Kar, A. Lemaçon, P. Soucy, D. Glubb, A. Rostamianfar, et al. Association analysis identifies 65 new breast cancer risk loci. *Nature*, 551(7678):92, 2017.

[19] A. P. Morris, B. F. Voight, T. M. Teslovich, T. Ferreira, A. V. Segre, V. Steinthorsdottir, R. J. Strawbridge, H. Khan, H. Grallert, A. Mahajan, et al. Large-scale association analysis provides insights into the genetic architecture and pathophysiology of type 2 diabetes. *Nature genetics*, 44(9):981, 2012.

[20] M. Nagel, P. R. Jansen, S. Stringer, K. Watanabe, C. A. de Leeuw, J. Bryois, J. E. Savage, A. R. Hammerschlag, N. G. Skene, A. B. Muñoz-Manchado, et al. Meta-analysis of genome-wide association studies for neuroticism in 449,484 individuals identifies novel genetic loci and pathways. *Nature genetics*, 50(7):920–927, 2018.

[21] C. P. Nelson, A. Goel, A. S. Butterworth, S. Kanoni, T. R. Webb, E. Marouli, L. Zeng, I. Ntalla, F. Y. Lai, J. C. Hopewell, et al. Association analyses based on false discovery rate implicate new loci for coronary artery disease. *Nature genetics*, 49(9):1385, 2017.

[22] C. M. Phelan, K. B. Kuchenbaecker, J. P. Tyrer, S. P. Kar, K. Lawrenson, S. J. Winham, J. Dennis, A. Pirie, M. J. Riggan, G. Chornokur, et al. Identification of 12 new susceptibility loci for different histotypes of epithelial ovarian cancer. *Nature genetics*, 49(5):680, 2017.

[23] B. P. Prins, K. B. Kuchenbaecker, Y. Bao, M. Smart, D. Zabaneh, G. Fatemifar, J. Luan, N. J. Wareham, R. A. Scott, J. R. Perry, et al. Genome-wide analysis of health-related biomarkers in the uk household longitudinal study reveals novel associations. *Scientific reports*, 7(1):1–9, 2017.

[24] J. C. Randall, T. W. Winkler, Z. Kutalik, S. I. Berndt, A. U. Jackson, K. L. Monda, T. O. Kilpeläinen, T. Esko, R. Mägi, S. Li, et al. Sex-stratified genome-wide association studies including 270,000 individuals show sexual dimorphism in genetic loci for anthropometric traits. *PLoS Genet*, 9(6):e1003500, 2013.

[25] S. Ripke, C. O'Dushlaine, K. Chambert, J. L. Moran, A. K. Kähler, S. Akterin, S. E. Bergen, A. L. Collins, J. J. Crowley, M. Fromer, et al. Genome-wide association analysis identifies 13 new risk loci for schizophrenia. *Nature genetics*, 45(10):1150, 2013.

[26] E. Sanderson, G. Davey Smith, F. Windmeijer, and J. Bowden. An examination of multivariable mendelian randomization in the single-sample and two-sample summary data settings. *International journal of epidemiology*, 48(3):713–727, 2019.

[27] J. E. Savage, P. R. Jansen, S. Stringer, K. Watanabe, J. Bryois, C. A. De Leeuw, M. Nagel, S. Awasthi, P. B. Barr, J. R. Coleman, et al. Genome-wide association meta-analysis in 269,867 individuals identifies new genetic and functional links to intelligence. *Nature genetics*, 50(7):912–919, 2018.

[28] F. R. Schumacher, A. A. Al Olama, S. I. Berndt, S. Benlloch, M. Ahmed, E. J. Saunders, T. Dadaev, D. Leongamornlert, E. Anokian, C. Cieza-Borrella, et al. Association analyses of more than 140,000 men identify 63 new prostate cancer susceptibility loci. *Nature genetics*, 50(7):928, 2018.

[29] H. Schunkert, I. R. König, S. Kathiresan, M. P. Reilly, T. L. Assimes, H. Holm, M. Preuss, A. F. Stewart, M. Barbalic, C. Gieger, et al. Large-scale association analysis identifies 13 new susceptibility loci for coronary artery disease. *Nature genetics*, 43(4):333–338, 2011.

[30] E. A. Stahl, G. Breen, A. J. Forstner, A. McQuillin, S. Ripke, V. Trubetskoy, M. Mattheisen, Y. Wang, J. R. Coleman, H. A. Gaspar, et al. Genome-wide association study identifies 30 loci associated with bipolar disorder. *Nature genetics*, 51(5):793–803, 2019.

[31] C. J. Willer, E. M. Schmidt, S. Sengupta, G. M. Peloso, S. Gustafsson, S. Kanoni, A. Ganna, J. Chen, M. L. Buchkovich, S. Mora, et al. Discovery and refinement of loci associated with lipid levels. *Nature genetics*, 45(11):1274, 2013.

[32] A. R. Wood, T. Esko, J. Yang, S. Vedantam, T. H. Pers, S. Gustafsson, A. Y. Chu, K. Estrada, Z. Kutalik, N. Amin, et al. Defining the role of common variation in the genomic and biological architecture of adult human height. *Nature genetics*, 46(11):1173, 2014.

[33] N. R. Wray, S. Ripke, M. Mattheisen, M. Trzaskowski, E. M. Byrne, A. Abdellaoui, M. J. Adams, E. Agerbo, T. M. Air, T. M. Andlauer, et al. Genome-wide association analyses identify 44 risk variants and refine the genetic architecture of major depression. *Nature genetics*, 50(5):668, 2018.

[34] M. Wuttke, Y. Li, M. Li, K. B. Sieber, M. F. Feitosa, M. Gorski, A. Tin, L. Wang, A. Y. Chu, A. Hoppmann, et al. A catalog of genetic loci associated with kidney function from analyses of a million individuals. *Nature genetics*, 51(6):957, 2019.

[35] C. X. Yap, J. Sidorenko, Y. Wu, K. E. Kemper, J. Yang, N. R. Wray, M. R. Robinson, and P. M. Visscher. Dissection of genetic variation and evidence for pleiotropy in male pattern baldness. *Nature communications*, 9(1):1–12, 2018.
